# Deciphering the Origin and Plasticity of Circulating Endothelial Cells: A Model for Systemic Angiogenesis Programs in Health and Disease

**DOI:** 10.1101/2025.08.26.672338

**Authors:** Orly Efros, Dror Brook, Yotam Yaniv, Jacob Elkahal, Nili Furer, Gal Dadi, Hadas Naor, Nimrod Rappoport, Oren Milman, Merav Kedmi, Uriel Katz, Sigal Tavor, Adrian Duek, Elad Maor, Leonid Sternik, Nathali Kaushansky, Eldad Tzahor, Karina Yaniv, Liran Shlush

**Affiliations:** Department of Molecular Cell Biology, Weizmann Institute of Science, Rehovot, Israel; National Hemophilia Center and Institute of Thrombosis & Hemostasis, Sheba Medical Center, Tel-Hashomer; Gray Faculty of Medical and Health Sciences, Tel Aviv-Yafo, Israel; Department of Computer Science and Applied Mathematics, Weizmann Institute of Science, Rehovot, Israel; Blavatnik School of Computer Science, Tel Aviv University, Tel Aviv, Israel; Department of Life Sciences Core Facilities, Weizmann Institute of Science, Rehovot, Israel; Pediatric Heart Institute, Edmond and Lily Safra Children’s Hospital, Sheba Medical Center, Tel-Hashomer, Israel; Hemato-Oncology Department, Assuta Medical Center, Tel Aviv, Israel; Maccabi Healthcare Services, Tel Aviv, Israel; Hematology Institute, University Hospital Samson Assuta Ashdod; Faculty of Health Sciences, Ben-Gurion University of the Negev, Beersheba, Israel; Leviev Cardiovascular Institute, Sheba Medical Center, Tel-Hashomer, Israel; Cardiac Surgery, Sheba Medical Center, Tel-Hashomer, Israel; Department of Immunology and Regenerative Biology, Weizmann Institute of Science, Rehovot, Israel; The Miriam and Aaron Gutwirth Medical School, Weizmann Institute of Science

## Abstract

Endothelial cells line the inner surface of blood and lymphatic vessels and play key roles in vascular function. Circulating endothelial cells (CECs) are endothelial cells found in the bloodstream, yet their origin and functional potential have not been fully elucidated.

To investigate their identity, we assembled and analyzed multiple public single-cell gene expression atlases comprising 212,144 endothelial cells from 23 human tissues. Using these datasets, we identified gene modules that quantitatively resolve endothelial types and states, including arterial, capillary, venous, lymphatic, and angiogenic-state programs, along with genes that are unique to endothelial cells of specific tissues.

Leveraging this knowledge to 597 CECs, which we isolated from 2.1 million circulating CD34⁺ cells across 287 donors, we found that CECs span a wide spectrum of mature endothelial identities and tissue-specific programs, consistent with diverse vascular origins. In culture, CECs downregulate mature endothelial markers (e.g., CD34) and acquire a progenitor-like PROCR⁺ transcriptional expression. Alteration of the Notch signaling *in-vitro* reverses this shift, with indirect activation of the pathway inducing a capillary and angiogenic program reminiscent of mature endothelial cells.

As we profiled the angiogenic program *in-vivo* and created an *in-vitro* angiogenic EC model, we sought to determine whether similar transcriptional programs are present in the context of cancer. We analyzed publicly available single-cell RNA-seq datasets from normal and malignant breast tissue, uncovering a marked enrichment of angiogenic endothelial cells not only within the tumor but, surprisingly, also in histologically normal contralateral breast tissue from breast cancer patients, pointing to a previously unrecognized systemic paraneoplastic effect. Transcriptomic overlap between these *in-vivo* angiogenic cells and Notch-biased cultured CEC derivatives suggests that CEC cultures may serve as an accessible human model for tumor-driven and paraneoplastic vascular remodeling.

Together, our study introduces a comprehensive framework for deciphering endothelial cell identity, illuminates CEC origin and plasticity, and provides a scalable platform to study cancer-associated endothelial reprogramming.

## Introduction

A complex network of interconnected blood and lymphatic vessels maintains organ health by delivering vital substances like oxygen and nutrients through a closed loop of arteries, veins, and capillaries while removing waste products. ^1^ At the core of this vascular system, endothelial cells (ECs) form the inner lining of these vessels, serving as critical gatekeepers. They regulate vascular tone, inflammatory response, maintain a non-thrombotic environment, and control the exchange of fluids, molecules, and circulating cells between the bloodstream and surrounding tissues. ^2–6^ Endothelial cells display significant phenotypic heterogeneity along the vascular tree, varying among distinct vascular types (e.g., lymphatic vessels, arteries, veins, and capillaries) and across tissues. ^1,7^ This diversity is evident in morphological differences, functional responses, and—as highlighted by integrated endothelial gene expression atlases in humans and murines—in distinct gene expression profiles that enable endothelial cells to execute specialized functions tailored to the unique requirements of various organs and physical conditions. ^8–12^ Notable examples include the blood-brain barrier, formed by brain endothelial cells to prevent neurotoxic plasma components, blood cells, and pathogens from entering the brain, as well as the role of liver sinusoidal cells in mediating the liver’s filtration and scavenger functions. ^9,13,14^

In addition to tissue-resident, a small population of endothelial cells circulates within the bloodstream. ^15^ First identified in the 1960s, the origin and functional role of circulating endothelial cells (CECs) remain largely unresolved. ^16,17^ The estimation of CEC remains controversial, as quantification reports rely on flow cytometry using various markers, some of which lack specificity for endothelial cells. ^17^ In healthy individuals, reported CEC counts range from a few to several thousand per milliliter, while in certain diseases, levels can rise to approximately 40,000 per milliliter. ^15,17,18^ Elevated levels have been associated with a variety of conditions, including cardiovascular diseases, solid and hematological malignancies, autoimmune disorders, and pathologies marked by mechanical vascular damage (e.g., sickle cell anemia). ^19–22^ These cells, which express *CD34* and can be cultured from peripheral blood, were shown to contribute to postnatal vasculogenesis and angiogenesis, leading to their designation as endothelial progenitor cells (EPCs). ^23–25^

In this study, we isolated CECs from over 2 million CD34-positive cells derived from the peripheral blood of 287 individuals, resulting in the capture of 597 cells. To elucidate their phenotype and potential roles, we constructed comprehensive single-cell transcriptomic models using publicly available datasets encompassing 23 tissues and 212,144 endothelial cells. Using these models, we validated vascular and tissue-specific markers and uncovered novel ones. Our analyses reveal that most CECs express markers characteristic of mature vascular types. After 3 and 6 weeks of culture, these cells demonstrated a further reduction in mature endothelial markers, including *CD34*, and an increase in *PROCR*, which has been implicated as a marker of endothelial stem cells. ^26^ Non-direct activation of the Notch signaling pathway in the cultured CECs reversed this phenotype, reinstating *CD34* expression along with additional capillary and angiogenic endothelial cell markers, while diminishing *PROCR* levels.

By analyzing single-cell transcriptomic datasets from normal, contralateral, and cancer breast tissues, we reveal an unexpected increase in angiogenic endothelial cell proportions in the contralateral breast tissue endothelial cells, indicating a paraneoplastic effect.

Collectively, we built comprehensive endothelial cell atlases spanning vascular types and organs across the adult human body. Using these atlases, we developed a computational tool for identifying endothelial types and states, which uncovered a novel systemic paraneoplastic angiogenic endothelial cells enrichment in breast cancer. By elucidating the origins of circulating endothelial cells and their *in-vitro* dedifferentiation and redifferentiation capacities, we establish a capillary angiogenic endothelial cell culture model to investigate the functional significance of this phenomenon.

## Results

### Quantified transcriptional characterization of endothelial type, state, and tissue identity

To identify the origin of circulating endothelial cells (CECs), we leveraged previous findings showing that endothelial cells exhibit distinct transcriptional signatures across vessel types (lymphatic, arterial, venous, and capillary) and organs and constructed a comprehensive reference map of human endothelial cells (**Fig. 1A**). ^9,11,12^

**Fig. 1:**
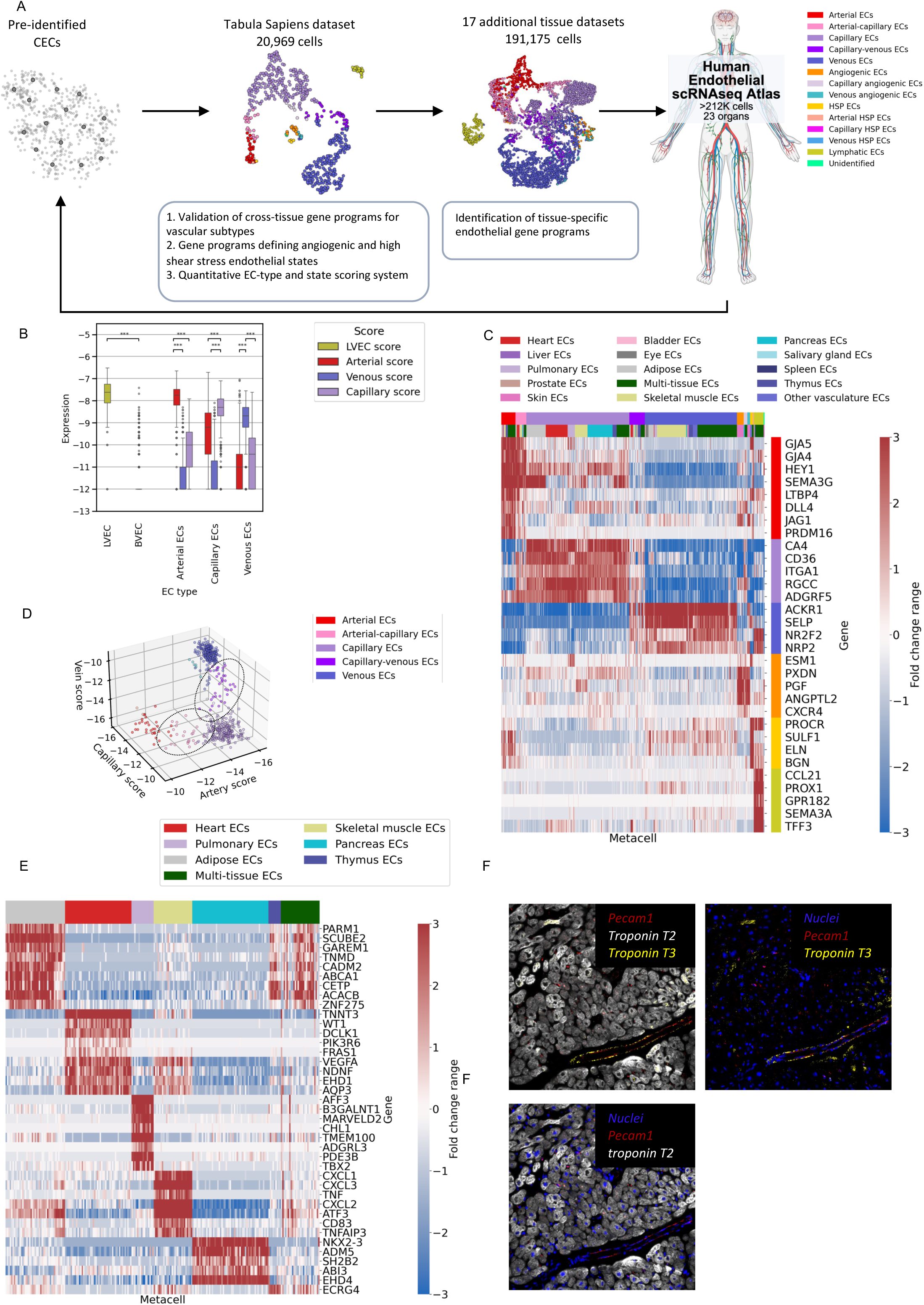
Quantified transcriptional characterization of human endothelial identity across types, states, and tissues. A. Workflow for mapping circulating endothelial cells (CECs) to their tissue of origin through construction of a comprehensive scRNAseq human endothelial cell (EC) reference atlas. Schematic representation showing a two-dimensional scRNAseq UMAP projection of the transcriptional manifold of (from left to right): CECs of unknown origin collected from 417 individuals; ECs from the Tabula Sapiens dataset; ECs from 17 healthy tissue datasets. Each dot represents a metacell, with colors representing EC type. These latter reference atlases, comprising 191,175 across 23 organs, enabled the development of a quantitative scoring framework for EC type, state, and tissue identity, into which the CEC model was projected to infer vascular type, state, and tissue of origin. Created with BioRender.com. B. Boxplots showing the log₂-transformed expression of endothelial subtype scores (lymphatic vessel endothelial cells [LVEC] versus blood vessel endothelial cells [BVEC], arterial, venous, capillary ECs) across individual cells, grouped by their original cell-type annotation. C. Heatmap showing literature-derived vascular endothelial type markers across the endothelial Tabula Sapiens metacell atlas. The top color bars indicate endothelial type (top bar; color as in Fig. 1A) and tissue of origin (bottom bar; additional color-code provided separately). The right color bar indicates marker type, with colors matching the corresponding cell type in Fig. 1A). D. 3D scatter plot of absolute BVEC subtype scores (arterial, venous, capillary) per metacell. Dashed circles highlight intermediate groups co-expressing two BVEC subtypes. E. Heatmap of differentially expressed, tissue-specific marker genes in capillary endothelial metacells from the endothelial Tabula Sapiens atlas. The top color bar indicates tissue of origin. F. Immunohistochemistry of human cardiac tissue. The image demonstrates the co-localization of Troponin T3 and the endothelial marker PECAM1 along the cardiac vasculature. Cardiomyocytes are stained for Troponin T2 (white), and nuclei with DAPI (blue).

First, we used the Tabula Sapiens, a comprehensive single-cell RNA-seq dataset spanning multiple human tissues and organ systems. ^27^ Endothelial cells (ECs) were extracted based on the known markers *CDH5* and *PECAM1* (Methods) (**Extended Data Fig. 1A**), resulting in a subset of 20,758 cells spanning 13 distinct tissues. To identify EC subpopulations, we constructed a metacell model, providing a robust quantitative representation of the transcriptional manifold (**Fig. 1A, Extended Data Fig. 1B**). ^28^

To annotate vascular type, we used the pre-annotated vascular type from the Tabula Sapiens dataset, where available, to evaluate established literature markers. Markers that distinguished one vascular type from others were grouped into gene modules. These gene modules provide a robust signal score for vascular types, which can be applied to classify endothelial types at the metacell and single-cell levels. Validated genes included, among others, *PROX1* and *SEMA3A* for lymphatic ECs; *DLL4*, *GJA4*, and *GJA5* for arterial ECs; *NR2F2* and *NRP2* for venous ECs; and *CA4* and *CD36* for capillary ECs (**Supplementary Table 1**). The lymphatic score effectively separates blood-endothelial cells (BVECs) from lymphatic endothelial cells (LVECs) (p-value < 0.05) and distinguishes BVEC cells into arterial, capillary, and venous subtypes (p < 0.05 for all pairwise comparisons) (**Fig. 1B, Extended Data Fig. 1C-F**).

Next, we annotated the entire endothelial manifold using these gene modules (**Fig. 1C**). While most metacells corresponded to a single vascular type, a subset displayed intermediate expression levels of two vascular scores. These included artery and capillary or capillary and vein, suggesting transcriptional profiles that bridge distinct vascular identities. Unsurprisingly, we found no transcriptomic state expressing both artery and vein signatures simultaneously.

To address metacells that did not match either vascular type, we extended our analysis to endothelial states that were not annotated in the original dataset. We constructed an angiogenic endothelial cell gene module using the previously established markers *ESM1*, *PXDN*, *PGF*, *ANGPTL2*, and *CXCR4*. ^29–32^ While some metacells only expressed angiogenic EC gene module, others co-expressed vascular type and angiogenic EC signatures, representing a mixed transcriptional state. This may hint at transitional states indicative of dynamic cellular processes, such as ongoing differentiation or plasticity between endothelial states. Finally, we annotated a subset of metacells, termed high-shear pressure endothelial cells (HSP ECs), expressing elevated levels of *SULF1*, *ELN*, *BGN*, and *PROCR*, genes that have been linked to ECs in large high-pressure vessels (**Extended Data Fig. 2A**). ^9,33^ Notably, *PROCR* has previously been implicated as a marker of endothelial progenitor cells. ^26,34^

Using this comprehensive type and state scoring system, we annotated all metacells for vessel type and state, except for two (**Fig. 1C**). The vascular scoring system showed effective discrimination between BVEC subtypes at the metacell level, while also capturing intermediate transcriptional states (**Fig. 1D**).

Using our annotated model, we identified highly expressed markers, previously unreported or described in specific tissues, for BVECs, including *IFI27*, and *SPARCL1*; for LVECs, including *MPP7* and *NR2F1*; and for BVEC subtypes, such as and *PCSK5* for arterial ECs, *PRCP* and *OLFM1* for venous ECs, and *BTNL9* and *TPM1* for capillary ECs (**Extended Data Fig. 2B).** These novel gene module scores effectively distinguish between BVEC and LVEC populations, and between BVEC subpopulations **(Extended Data Fig. 2B**). To note, *CD34* exhibited distinct increased expression among BVECs, while its expression was markedly low in LVECs and across most angiogenic ECs. ^35,36^ Some venous and lymphatic markers, including *NR2F2* and *NRP2*, showed overlapping patterns of high expression.^37^

To identify tissue-specific markers, we analyzed the Tabula Sapiens endothelial model by tissue of origin. Capillary endothelial metacells predominantly mapped to a single tissue of origin. In contrast, a substantial proportion of arterial and venous metacells reflected mixed tissue origins (51.5% in arterial ECs and 36% in venous compared to 10.8% in capillary ECs; **Extended Data Fig. 2C**), suggesting greater transcriptomic similarity across tissues in arterial and venous endothelial cells than in capillary cells.

Tissue-specific transcriptomic analysis of endothelial cells uncovers distinct, organ-specific expression patterns (**Fig. 1E, Extended Data Fig. 3A, B**). Throughout the vascular endothelial cell types, thymic ECs exhibited elevated levels of adipose markers, such as *CETP* and *ACACB*, which may reflect age-related thymic involution and the associated accumulation of adipose tissue. ^38^ Similarly, a substantial gene overlap was observed between capillary ECs of cardiac and skeletal muscle, including genes such as *VEGFA*, *EHD1*, and *AQP3*, reflecting their shared muscle tissue characteristics.

Interestingly, *TNNT3*, which encodes the fast skeletal muscle isoform, was highly expressed only in cardiac capillary and arterial endothelial cells (**Fig. 1E, Extended Data Fig. 3A**). ^39^ This expression pattern was confirmed in a cardiac endothelial cell model derived from an additional independent dataset (**Extended Data Fig. 3C**). When examining the expression of *TNNT3* in the full Tabula Sapiens atlas, its levels were low in cardiomyocytes yet high in cardiac endothelial cells, comparable to those observed in fast skeletal muscle (**Extended Data Fig. 3D)**. We further validated the protein-level expression of the protein Troponin T3 in cardiac ECs by immunofluorescence staining on fixed cardiac sections using a TNNT3-specific primary antibody. TNNT3 exhibited co-localization with endothelial markers, displaying a linear vascular pattern, while showing minimal to no signal in surrounding cardiomyocytes (**Fig. 1F**).

To expand the representation of ECs across human tissue and organs, we analyzed 17 publicly available scRNA-seq datasets spanning 15 tissues and comprising a total of 191,175 additional endothelial cells (**Supplementary Table 2**). We built a metacell model for each dataset and annotated them using the vascular-type scoring system described earlier (**Fig. 1A, Extended Data Fig. 3E)**. While each gene module consistently distinguishes vascular type across tissue models, the expression levels of individual markers vary depending on the tissue of origin (**Extended Data Fig. 3F**). Accordingly, the overall vascular-type scores display varying expression levels across tissues when compared to the Tabula Sapiens dataset (**Extended Data Fig. 4A**). Our novel identified markers effectively separate vascular subtypes, even in tissues not represented in the Tabula Sapiens model

Analysis of the additional datasets enabled a comprehensive view of organotypic specialization across endothelial cell types (**Extended Data Fig. 4B, C**). Studying genes enriched in distinct vascular endothelial subtypes and tissues may help uncover connections between their expression patterns and vascular diseases associated with those specific endothelial populations. Some examples include *BTNL9*, a gene previously shown to be increased in placental microvascular endothelial cells from severe IUGR cases, which was found in our analysis to be distinctly enriched in capillary ECs, hinting at a link to endothelial dysfunction in IUGR. ^40^ The *PLAU* gene, which was previously implicated in coronary atherosclerosis development, was enriched in cardiac arterial endothelial cells (**Extended Data Fig. 3A**). ^41^

Overall, we constructed a comprehensive endothelial cell atlas encompassing a wide range of tissues across the body. Our robust vascular-type scoring system effectively distinguishes between key endothelial subtypes—such as arterial, venous, capillary, and lymphatic populations—and various intermediate or distinct functional phenotypes (angiogenic ECs and HSP ECs). Furthermore, our models capture the tissue-specific transcriptional signatures within these vascular endothelial subtypes.

### Circulating ECs originate from mature endothelial cells of various types and tissues

As part of an atlas of circulating CD34+ (cCD34+) cells developed in our lab, peripheral blood samples were collected from 287 adult volunteers, yielding an enrichment of 2,053,909 cCD34+ cells. Among these, 121 samples were obtained from healthy individuals, while 166 were obtained from patients with hematological malignancies, including myelodysplastic syndrome (MDS) and myeloproliferative neoplasms (MPN) (**Fig. 2A**). ^42^ A subset of cCD34+ cells exhibited high expression of *CDH5* and *PECAM1* and were therefore identified as circulating endothelial cells **(Fig. 2B)**. These circulating endothelial cells (CECs) were unified into a single metacell model comprising 597 cells, representing 88 healthy individuals and 93 patients with hematological malignancies (**Supplementary Table 3, Fig. 2C)**. This number suggests that the number of CEC is lower than previously reported.^43^ The extreme rarity of CECs precluded follow-up analyses and enrichment from individual samples.

**Fig. 2:**
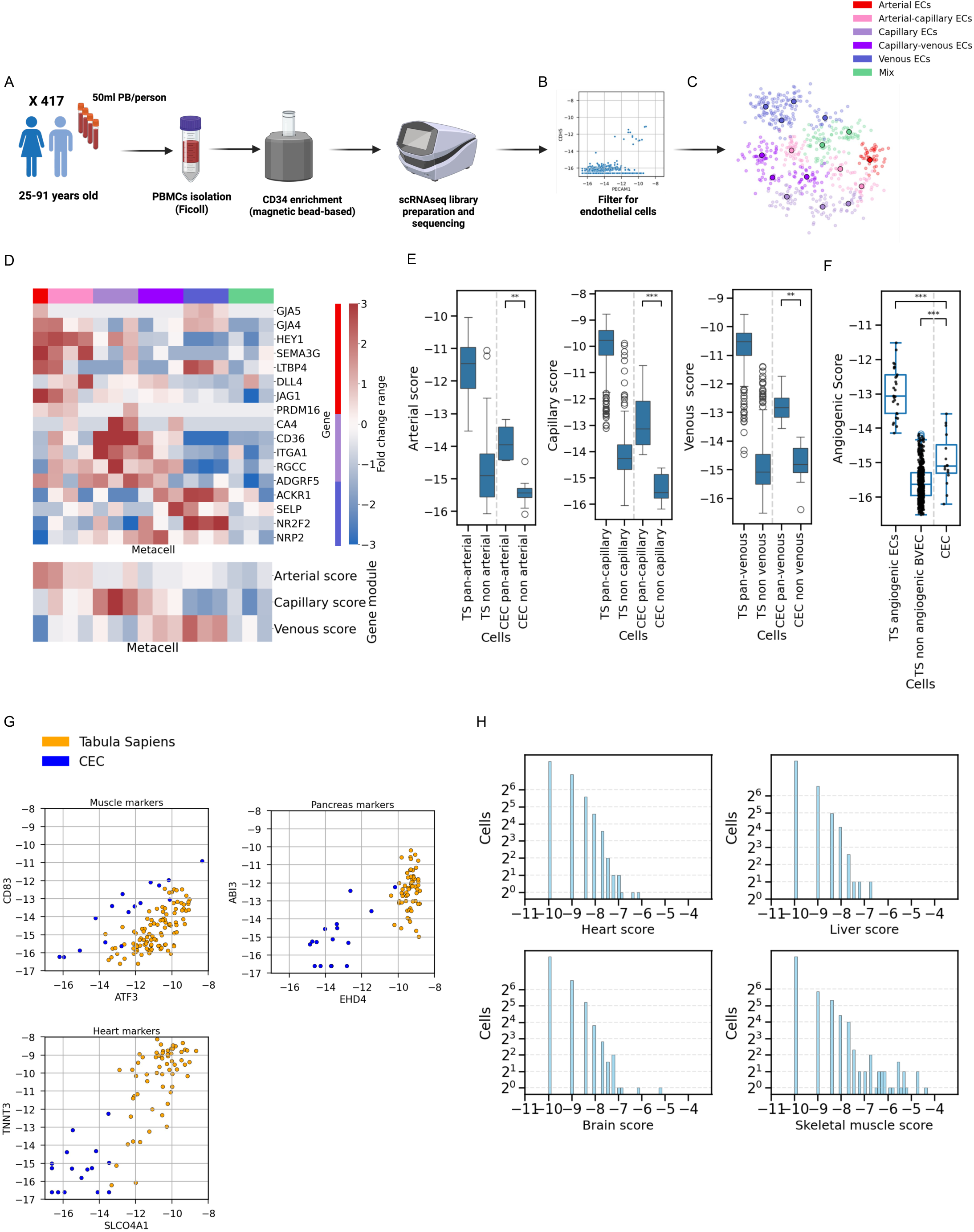
Circulating ECs originate from mature endothelial cells of various types and tissues. A. Experimental workflow for creating a scRNA-seq atlas of circulating endothelial cells (CECs). Created with BioRender.com. B. Scatter plot showing expression of canonical endothelial markers *PECAM1* and *CDH5* across the circulating CD34⁺ metacell manifold. A small subset of cells co-expresses both markers, consistent with circulating endothelial cell identity. C. Two-dimensional UMAP projection of the transcriptional manifold of 597 CECs. Colors indicate the annotated state of each metacell, with small dots representing individual cells assigned to that metacell. D. Heatmap of literature-based vascular type endothelial markers and computed scores across circulating endothelial metacells. The top color bar indicates the annotated endothelial type. E. Boxplots of arterial (left), capillary (center), and venous (right) blood vessel endothelial cells subtype scores in the endothelial Tabula Sapiens (TS) reference atlas and CECs. Each comparison shows subtype-positive and subtype-negative cells within both datasets, showing that CECs consistently exhibit a lower mature vascular subtype relative to tissue-resident endothelial cells. F. Boxplot of angiogenic scores across three groups of endothelial metacells: Tabula Sapiens (TS) angiogenic endothelial cells, TS blood vessel endothelial cells (BVECs), and CECs. CEC metacells show a higher angiogenic score compared to non-angiogenic BVECs and lower than angiogenic ECs. G. Scatter plots of tissue-specific marker gene expression in circulating endothelial metacells (blue) and tissue-matched endothelial metacells from the endothelial Tabula Sapiens reference atlas (orange). Each dot represents a metacell. While circulating endothelial metacells show lower expression of tissue markers, some retain detectable expression, suggesting a possible origin from corresponding mature tissues. The panel shows markers for (clockwise): muscle, pancreas, and heart. H. Histograms of tissue-specific score expression (heart, liver, brain, muscle, and skeletal muscle) across individual circulating endothelial cells, downsampled to 2¹¹ UMIs per cell. While most cells show low expression, a subset displays elevated tissue-specific scores, suggesting a potential tissue-of-origin signature.

These endothelial metacells exhibited low LVEC and high BVEC scores, comparable to the BVEC score levels observed in blood vascular endothelial metacells from the Tabula Sapiens model, suggesting a blood vasculature origin (**Extended Data Fig. 5A**). However, since the enrichment was CD34-based and LVECs lack *CD34* expression, we cannot rule out the presence of these cells in circulation.

We annotated the metacells using the vascular type gene modules described earlier. Consistent with the endothelial Tabula Sapiens model, CECs encompassed the full range of endothelial subtypes, including arterial, venous, capillary, and intermediate populations with combined arterial-capillary and capillary-venous characteristics **(Fig. 2D)**. This diversity suggests that these cells originate from the shedding of mature blood vascular endothelial populations. This also excludes the possibility that endothelial cells from the punctured vein were the sole source of the analyzed cells. Three metacells, representing 109 cells, while showing comparable *PECAM1* and *CDH5* levels with other metacells, had no clear vascular-type transcriptomic signature. Two of these unidentified metacells expressed high levels of the injury-related gene *S100A10* **(Extended Data Fig. 5B)**, implying a possible injury response rather than a defined endothelial lineage.

While CEC could be classified into different vascular types based on the vascular gene modules, the absolute expression of individual marker genes and the overall expression of vascular gene modules were decreased compared to the mature expression in the Tabula Sapiens dataset (**Fig 2E**). This decrease in expression, while keeping a comparable BVEC score, suggests that gene regulation related to vascular type markers may be affected by their specific microenvironment and is downregulated when the cells exit their native niche.

CECs express angiogenic gene signatures at higher levels than non-angiogenic ECs in the Tabula Sapiens EC model, yet lower than angiogenic ECs (**Fig. 2F**). Additionally, angiogenic ECs in the Tabula Sapiens EC model show lower levels of *CD34* compared to mature arterial, venous, or capillary ECs. While this could have reduced their recovery with a CD34-based enrichment, their signal still exceeds the capture threshold of our circulating-CD34 protocol, as seen in the circulating hematopoietic stem cells atlas (**Extended Data Fig. 5C**). Together with the fact that angiogenic ECs comprise only about 4.4% of Tabula Sapiens ECs, only a minute fraction is expected to appear in peripheral blood. When comparing the HSP score, CEC metacells in the CEC atlas exhibited a low HSP score (**Extended Data Fig. 5D**).

Cell cycle analysis showed two metacells with elevated expression of M-phase–associated genes, including *MKI67* and members of the *KIF* gene family, indicative of active cell cycle progression (**Extended Data Fig. 5E**). ^44,45^ The levels of this gene module expression were higher compared to the ECs in our Tabula Sapiens-based EC atlas. These metacells included cells from healthy individuals and patients with hematological malignancies (44.9% and 58.8%, respectively). This may reflect endothelial cells undergoing division at the time of niche exit, or, less likely, continued proliferation while in circulation.

Tissue-specific analysis revealed metacells expressing gene signatures characteristic of muscle, heart, pancreas, and other organ-specific endothelial populations (**Fig. 2G**). Their absolute expression levels were lower than their mature model counterparts, mirroring the decrease in vascular type markers. Given the small sample size, each metacell likely aggregates cells from multiple tissues, which may also contribute to the overall reduction in tissue-specific marker expression. To improve tissue-level resolution and minimize cross-tissue signal overlap, we performed a secondary analysis, focused on tissues exhibiting robust tissue-specific differentially expressed signals detectable at the single-cell level (**Fig. 2H, Extended Data Fig. 5F)**. As expected, given the small sample size, most cells could not be assigned to a specific tissue. Nonetheless, 34 cells showed high expression levels of gene modules matching the heart, liver, brain, muscle, kidney, and lymph node. Crucially, neither analysis revealed a bone-marrow– specific signature. Together, these two complementary approaches provide the first evidence that CECs arise from diverse tissues.

Collectively, CECs show transcriptional heterogeneity indicative of origins from diverse blood vascular subtypes and organs despite reduced expression of subtype-specific and tissue-specific markers, likely due to their displacement from native tissue environments.

### Circulating endothelial cells undergo dedifferentiation in vitro

To further characterize CECs and to examine whether they maintain their mature EC markers *in-vitro*, we cultured them and analyzed them using scRNA-seq. Peripheral blood mononuclear cells (PBMCs) of 13 healthy individuals (age range 26-68) were cultured under endothelial-specific conditions, as described in previous papers (**Fig. 3A**). ^46^ We identified cells with the characteristic morphology of endothelial cells after a minimum of ten days in culture, with those cells becoming the majority of cells around three weeks **(Fig. 3B)**. When passed from their primary culture to a Matrigel surface, these cells spontaneously formed tubes, characteristic of endothelial cells (**Fig. 3C)**. ^47^

**Fig. 3:**
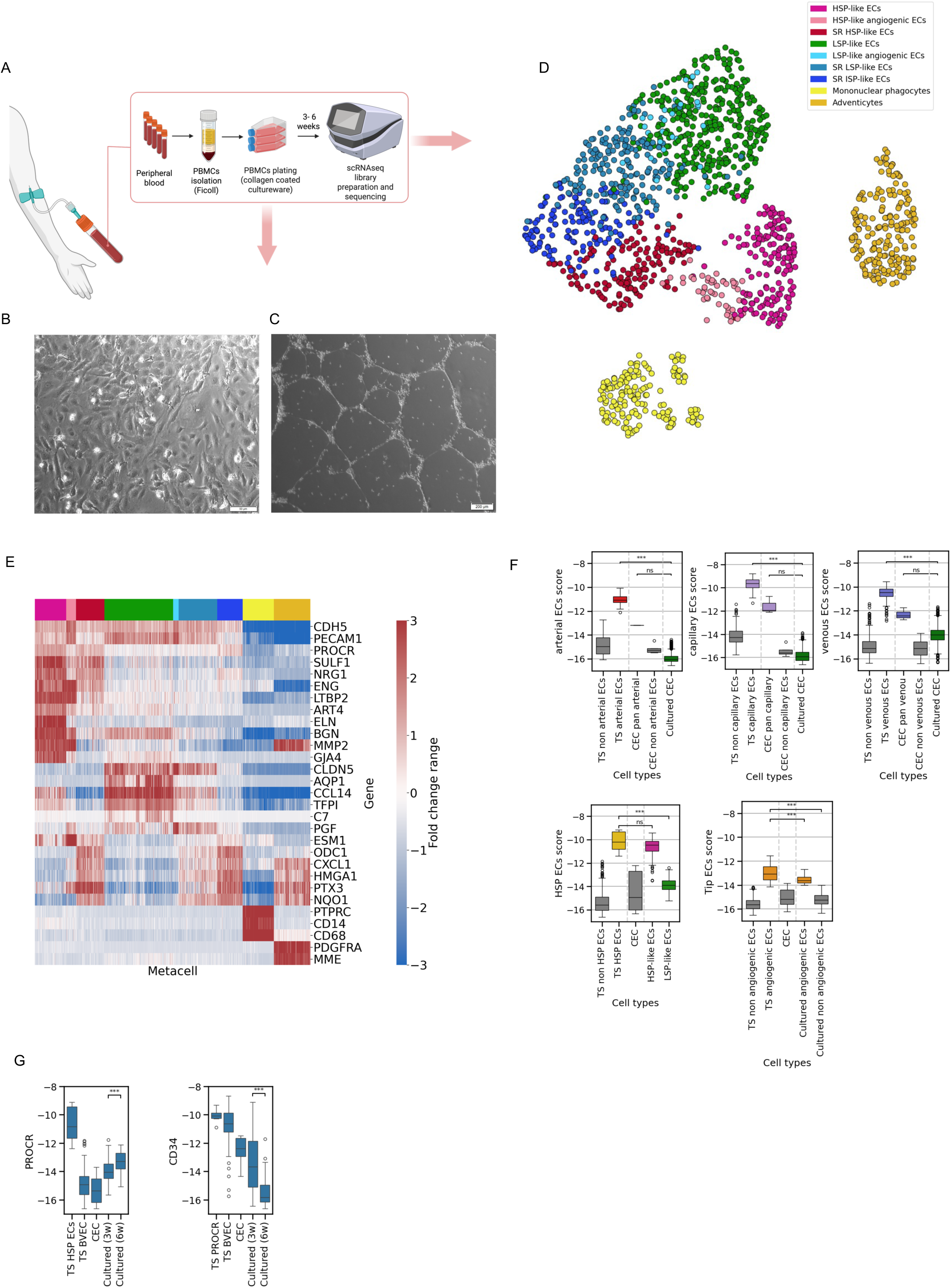
Circulating endothelial cells undergo dedifferentiation in vitro. A. Experimental workflow for creating a scRNA-seq model of cultured circulating endothelial cells (CECs). Created with BioRender.com. B. Phase-contrast microscopy images of cultured CECs after three weeks, showing the characteristic cobblestone morphology of endothelial monolayers. C. Phase-contrast microscopy images of cultured CECs in a Matrigel tube formation assay, CECs organize into capillary-like network structures within six hours of incubation. D. Two-dimensional UMAP projection of the transcriptional manifold of the cultured circulating endothelial cells model, with colors representing the annotated states of each metacell. E. Heatmap of differentially expressed genes across cultured CEC metacells. The top color bar indicates the annotated metacell state. F. Boxplots of arterial, capillary, venous, lymphatic, and angiogenic endothelial subtype scores across metacells from Tabula Sapiens (TS), CEC, and cultured CEC datasets. The boxplot color corresponds to the endothelial type shown in Fig. 3D. G. Boxplots of *PROCR* (left) and *CD34* (right) expression levels across endothelial cells from the endothelial Tabula Sapiens (TS) reference atlas, CEC, and cultured CECs at 3 and 6 weeks, showing an increase in *PROCR* and a decrease in *CD34* expression during culture.

We collected and performed scRNA-seq on cultured cells after three and six weeks post-seeding. After quality control and filtering, we retained 65,577 single-cell profiles and created a metacell model (**Fig. 3D**). We annotated metacells with high expression of *CDH5* and *PECAM1* as endothelial metacells. We also found mononuclear phagocyte cells, characterized by the expression of markers such as *PTPRC*, *CD14*, and *CD68*, both in the three and six-week cultures^48^. In a single culture, a population expressing *PDGFRA* and *MME*, consistent with perivascular adventitial cells, was detected, likely resulting from inadvertent sampling during vein puncture. ^49,50^

The endothelial cell population comprised five main subpopulations, with two additional subsets showing angiogenic-EC state signature, according to our previously described gene modules, stemming from these groups **(Fig. 3E)**. These cultured cell phenotypes may be induced by cultured mechanical and contact pressure and stress as previously described. One group of ECs showed similarity to the HSP-ECs seen in the Tabula Sapiens model, with elevated expression of genes associated with high-shear pressure exposure, including *PROCR*, *SULF1*, *ELN*, and *BGN* (**Fig. 3F, Extended Data Fig. 6A**). ^9,33^ This subset also exhibited high levels of additional genes shared with the HSP-ECs, such as extracellular matrix components and remodeling factors (*LTBP2, MMP2*), as well as *NRG1*, *ART4*, and arterial markers like *GJA4* and *LTBP4*. ^51–54^ Therefore, we named this group high-shear pressure–like endothelial cells (HSP-like ECs), with a subset exhibiting elevated angiogenic-EC scores referred to as HSP-like angiogenic-ECs.

Another group, composed of a subset of ECs with elevated angiogenic-EC scores, lacked high expression of genes characteristic of HSP-like ECs and had strong expression of genes enriched in venous endothelial cells from the Tabula Sapiens atlas. This group was designated low-shear pressure-like ECs (LSP-like ECs). These genes included *AQP1, C7, TFPI*, and *CCL14*. We speculated that, consistent with their venous similarity, typically associated with lower shear stress and slower blood flow, these cells are functionally linked to vascular homeostasis. For example, *TFPI* is a key inhibitor of the coagulation cascade, supporting an anti-thrombotic endothelial surface; *AQP1* encodes a water channel involved in fluid balance, and *C7* has been implicated in regulating endothelial inflammatory responses. ^55–57^

Three endothelial subpopulations were characterized by stress-associated gene expression, including *NQO1* and *HMGA1*, and exhibited varying levels of markers typical of HSP-like and LSP-like ECs, and were named accordingly (i.e, stress-related [SR] HSP-like, SR LSP-like, and SR intermediate shear pressure [ISP] ECs). ^58,59^

Cultured CEC subpopulations consistently exhibited low scores for mature vascular types, including BVEC, LVEC, arterial, capillary, and venous ECs, compared to Tabula Sapiens ECs and to the CECs (**Fig. 3F**). This suggests that CECs lose mature vascular signatures during *in vitro* culture.

After 3 weeks in culture, endothelial cells showed variable expression levels of the BVEC marker *CD34*, which declined by 6 weeks. In contrast, *PROCR* expression increased over time in culture, inversely correlated with *CD34* levels (Spearman’s ρ = –0.673, p < 0.05). When compared to the ECs in the Tabula Sapiens model and fresh CECs, we see that *CD34* expression steadily decreases from *in vivo* ECs (i.e., Tabula Sapiens dataset) to CECs, to 3-week cultures, and declines further at 6 weeks, with an opposing trend in *PROCR* expression (**Fig. 3G**). This dynamic demonstrates the plasticity potential of these cells and suggests that while CD34 is a marker of mature ECs, increased *PROCR* expression in culture may represent a loss of mature vascular type phenotype. Overall, our findings reveal marked phenotypic plasticity in circulating endothelial cells, characterized by the loss of mature vascular markers and acquisition of progenitor-like features during *in vitro* culture.

### Cultured circulating endothelial cells re-differentiate in response to notch signaling alteration

To test the potential of cultured CEC to re-differentiate, we partially adopted a differentiation protocol for arterial and venous endothelial cells from pluripotent stem cells, expected to non-directly activate and inhibit the Notch signaling pathway. ^60,61^ Twelve hours after applying the different media on the cultured circulating endothelial cells, we observed distinct morphological changes under light microscopy in the non-direct Notch pathway activation (**Fig. 4A, Extended Data Fig. 7A**). Notably, cells in the Notch activation condition displayed prominent elongation across the entire well. The morphological changes in the Notch inhibition condition were less pronounced compared to the control condition cells (**Fig. 4B**).

**Fig. 4:**
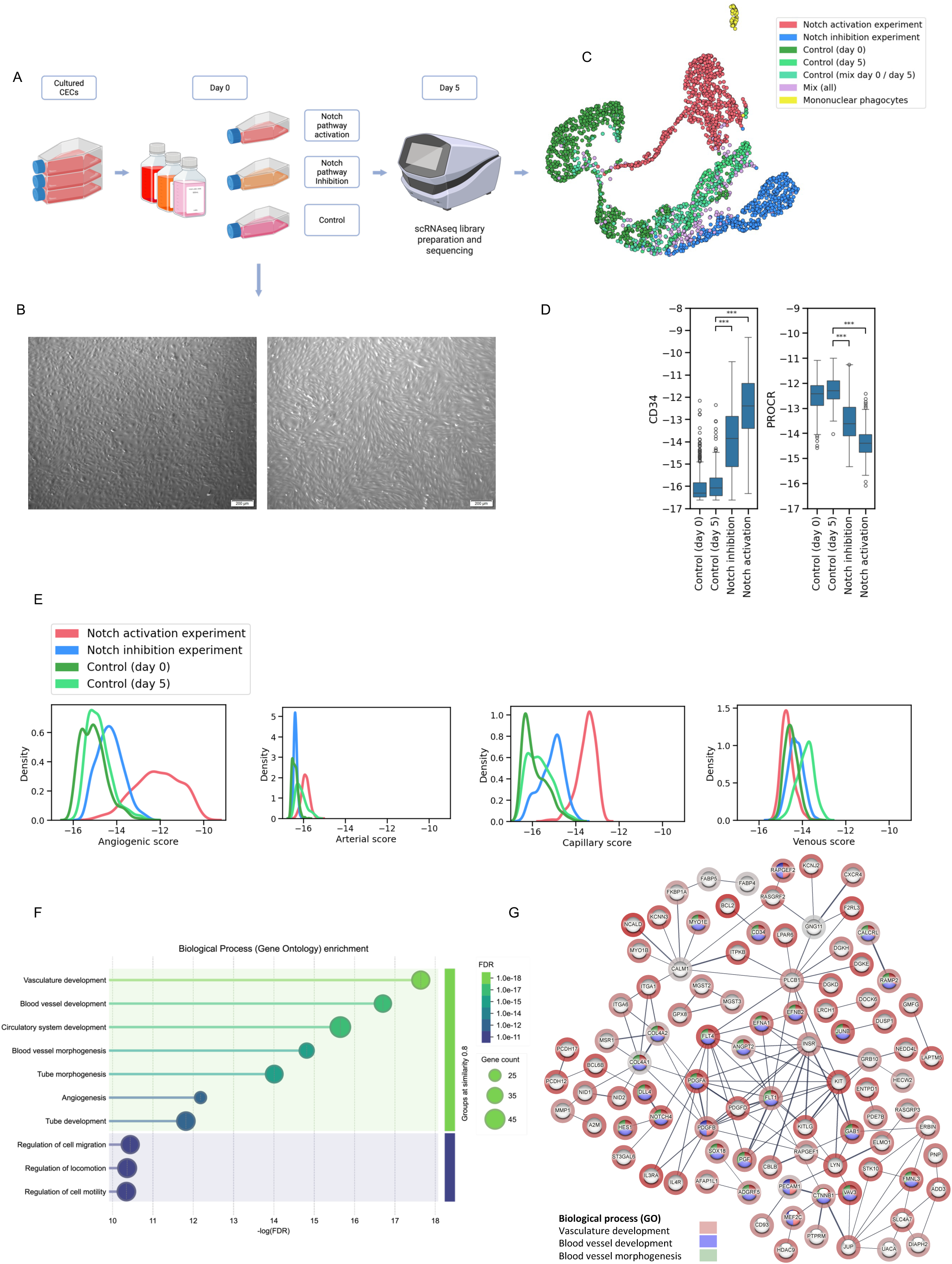
Cultured circulating endothelial cells re-differentiate in response to notch signaling alteration. A. Experimental workflow for creating a scRNA-seq model of cultured CECs following Notch signaling pathway activation and inhibition. Created with BioRender.com. B. Phase-contrast microscopy images of cultured CECs. Notch signaling activation (right) showing elongated morphological changes compared to standard adherent culture conditions (left). C. Two-dimensional UMAP projection of cultured CEC transcriptional manifold following notch signaling alteration, with colors representing annotated metacell state. D. Boxplots of *CD34* (left) and *PROCR* (right) expression levels in cultured circulating endothelial cells under control conditions (day 0 and day 5), Notch inhibition, and Notch activation. Notch signaling alteration reverses the previously observed trends, with *CD34* expression increasing and *PROCR* expression decreasing. E. Density plots showing angiogenic and blood vascular endothelial cell (arterial, capillary, venous) subtype scores in cultured circulating endothelial cells under different conditions. Notch activation significantly increases angiogenic and capillary scores compared to control. F. Gene Ontology enrichment analysis of Notch-activated circulating endothelial cells. The top ten enrichment programs, ranked by FDR significance, are shown. The analysis was based on differentially expressed genes (Notch activation vs. control) and on the top ten correlated genes from the 20 strongest DE genes (duplicates removed). Results were obtained using STRING v11.5 (Szklarczyk et al., 2021; https://string-db.org). G. String network analysis of Notch-activated circulating endothelial cells. The analysis was based on differentially expressed genes (Notch activation vs. control) and on the top ten correlated genes from the 20 strongest DE genes (duplicates removed). Interaction evidence was derived from experiments, databases, co-expression, neighborhood, gene fusion, and co-occurrence (minimum score 0.4). Disconnected nodes were omitted; node halos indicate gene expression values, and edge thickness reflects interaction support. Analysis performed with STRING v11.5 (Szklarczyk et al., 2021; https://string-db.org).

After five days, cells from each condition were subjected to scRNA sequencing, generating 114,532 cells that were integrated into a metacell model. Endothelial metacells were annotated based on their condition, using a majority vote of the cells in each metacell (i.e., >75%) (**Fig. 4C**). Most metacells represented a single condition or a mixture of control condition cells (day 0 and day 5).

As expected, following any of the Notch signaling pathway alterations, the gene expression of *CD34* increased, while the *PROCR* levels decreased, reversing the process we observed in the cultured cells without such signals (**Fig. 4D**).

In the Notch activation condition, an upregulation of the arterial EC marker DLL4, NOTCH4, and JAG2 was observed, as well as ARL15, which our endothelial-type scoring system classified as an arterial marker (**Extended Data Fig. 7A**). In addition, the Notch activation condition showed increased expression of angiogenic endothelial cell markers (*CXCR4, ESM1, INSR, PLVAP*, and *PXDN*) and capillary endothelial markers (*ADGRF5, ITGA1*, and *RGCC)* (**Supplementary Table 4, Extended Data Fig. 7B**). ^11,29,62,63^ Notably, *LYVE1*, a lymphatic endothelial cell marker, was enriched under Notch-inhibitory conditions, contrasting an earlier report in which blocking the Notch ligand *DLL4* reduced *LYVE1* expression in cultured lymphatic ECs. ^64^ This may indicate that broader Notch pathway inhibition differentially influences lymphatic marker expression.

Consistent with these findings, metacells enriched in Notch activation condition cells displayed prominently elevated angiogenic-EC and capillary scores, along with reduced venous scores (**Fig. 4E**). Specifically, these cells exhibited a capillary endothelial score comparable to that of mature capillary endothelial cells in both the Tabula Sapiens and individual tissue models, demonstrating their ability to re-differentiate in response to external signals. The Notch inhibition condition did not increase venous scores compared to control cells, suggesting that differentiation towards venous may require additional signals beyond Notch pathway suppression (**Fig. 4E**).

Gene ontology enrichment analysis of the Notch activation condition demonstrated a significant enrichment of vessel development–related programs (**Fig. 4F**). ^65^ STRING analysis of the Notch activation condition compared to the control further identified upregulation of genes associated with the Notch signaling pathway, including *NOTCH4, DLL4*, and *HES1* (**Fig. 4G**). ^65–67^

Together, our findings underscore the dynamic responsiveness of cultured circulating endothelial cells to Notch signaling pathway modulation, unveiling their capacity for transcriptomic reshaping under specific signaling conditions.

### Paraneoplastic angiogenic endothelial cell transformation in breast cancer

Having mapped the angiogenic transcriptional program *in vivo* and established an *in vitro* model of angiogenic endothelial cells, and given previous reports that cancer-associated ECs often display a tip-like phenotype, we sought to quantify this phenotype in cancer tissue. ^63^ We therefore constructed a metacell model of ECs from a publicly available dataset of breast cancer and normal tissue. ^68^ The model was annotated using our scoring system (**Fig. 5A, left**). As expected, after filtration for samples containing more than 50 ECs, a significantly larger fraction of ECs from cancerous breast tissue was classified as angiogenic ECs compared to the normal tissue (median 32.7% vs 3.5% respectively) (**Fig. 5A, right; Fig. 5B**). This trend was consistent across breast cancer subtypes, including ER+, HER2+, PR+, and triple-negative breast cancers (**Fig. 5C**). We found no significant difference between the BRCA1 and normal patients (**Fig. 5C**). At the individual patient level, we observe various levels of angiogenic EC proportion, both in the healthy samples and cancer (**Extended Data Fig. 8A**).

**Fig. 5:**
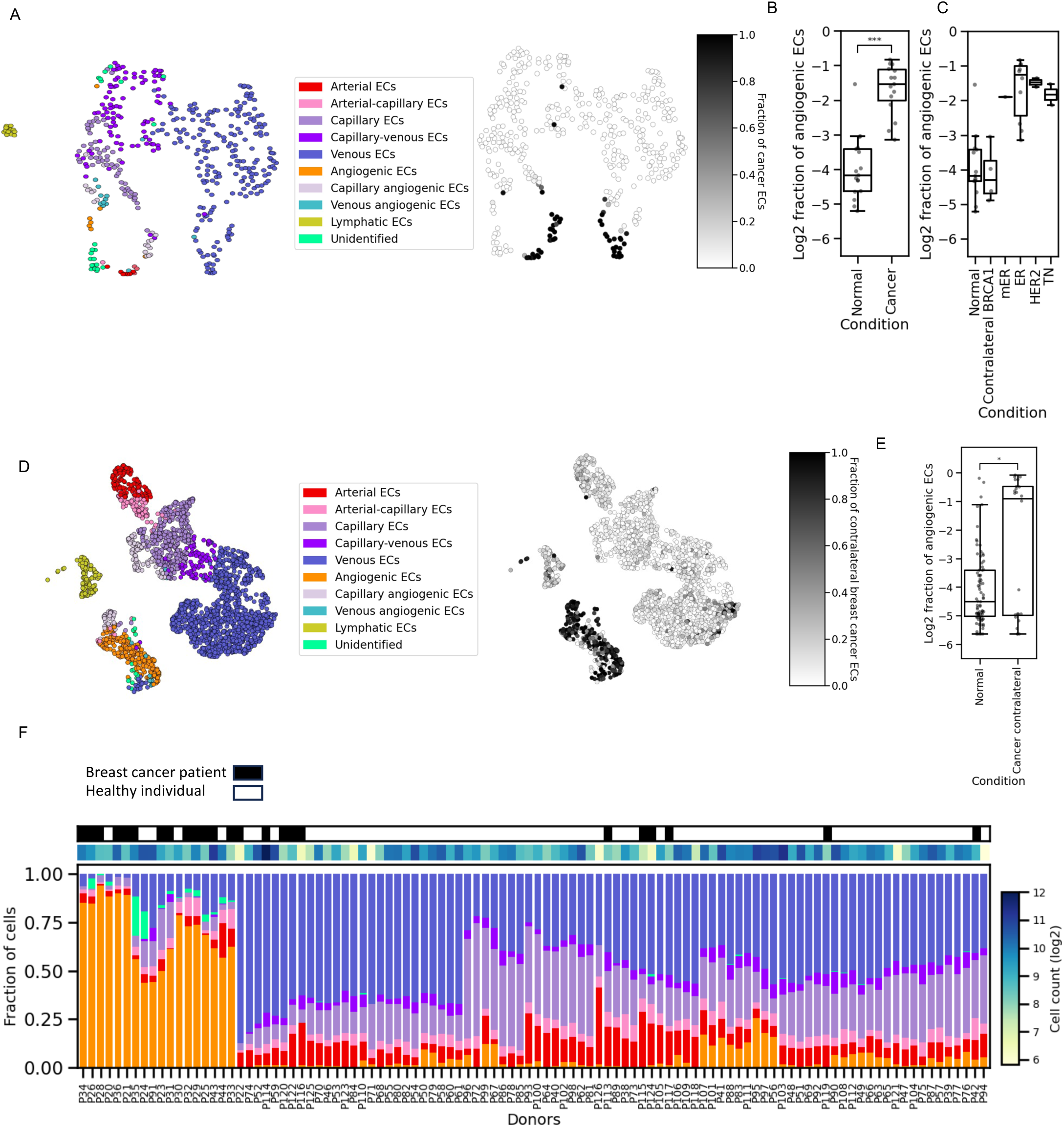
Paraneoplastic systemic angiogenic endothelial cell transformation in breast cancer. A. Two-dimensional UMAP projection of the transcriptional manifold of endothelial cells from the Chen, Y., Pal, B., Lindeman, G.J. *et al*. breast cancer dataset, showing metacells colored by type and state (left) and by the fraction of cells originating from breast cancer tissue. B. Boxplots of angiogenic cell fraction distributions across individuals in normal breast tissue versus breast cancer tissues (log scale). Only patients with at least 50 cells were included. C. Boxplots of angiogenic cell fraction distributions across individuals in normal breast tissue versus breast cancer tissues, with separation to cancer type (log scale). Only patients with at least 50 cells were included. D. Two-dimensional UMAP projection of the transcriptional manifold of endothelial cells from the Kumar *et al.* breast cancer dataset, showing metacells colored by annotated endothelial state (left) and by the fraction originating from contralateral non-tumor tissue (right). E. Boxplots of angiogenic cell fraction distributions across normal breast tissue versus contralateral breast cancer tissue (log scale). Only patients with at least 50 cells were included. F. Endothelial cell state composition per patient. Each vertical bar represents one patient, with colors indicating proportions of annotated cell states. Only patients with at least 50 cells are shown.

Surprisingly, during the analysis of the publicly available normal breast endothelial cell data from *Kumar et al.*, we found an unusually high proportion of endothelial cells in an angiogenic EC state in the contralateral breast tissue of breast cancer patients. ^69^ To address this, we constructed two models from this dataset: one comprising only healthy donor breast tissue samples (reduction mammoplasty and prophylactic mastectomy), which was included in the individual tissue model of healthy breast tissue (**Fig. 1**), and a second that included contralateral mastectomy tissue from breast cancer patients (healthy donors: n=81, with 55,817 ECs, contralateral mastectomy: n=26, with 15,909 ECs) (**Fig. 5D**, **left**). Samples containing fewer than 50 endothelial cells were excluded to ensure adequate representation, which resulted in the removal of one healthy individual sample and two cancer patient samples. Similar to other tissue models, we annotated the full model using the vascular-type gene modules and found a high representation of angiogenic ECs **(Fig. 5D)**. While a subset of angiogenic ECs co-express vascular-type gene modules, the majority do not show a vascular-type identity. Further analysis showed that a large portion of these cells was contributed from the contralateral tissue samples (**Fig. 5E**). Specifically, healthy individual samples had a median of 2.4% of angiogenic ECs, whereas contralateral breast cancer samples had a median of 51.8%, displaying a clear bimodal distribution (**Fig. 5C, Extended Data Fig. 8B**). To further analyze these individual differences, we performed a cell state composition analysis at the patient level (**Fig. 5F).** We found that most healthy individual samples had a low fraction of angiogenic ECs, while a few had a comparably high proportion of angiogenic ECs as the contralateral tissue. Conversely, most contralateral breast samples contained a large proportion of angiogenic ECs, while some displayed low levels that were similar to those observed in the healthy individual samples. The healthy individual samples were categorized into three groups based on their angiogenic EC fraction: the first, comprising 51/80 individuals (63.7%), showed low angiogenic EC levels (<5%). The second group included 24 individuals (30%) with intermediate angiogenic EC levels (range 5% - 25.2%). The third group, consisting of 5 individuals (6.25%), had high levels of angiogenic ECs (range 46%-87.5%). The contralateral breast cancer patient samples had a bi-modal distribution with 13/24 (54%) of patients exhibiting a high proportion of angiogenic ECs (range 48.6% - 92.5%), while the others (46%) had low levels (<3.8%). These results remained consistent after filtering for samples containing more than 500 cells (**Extended Data Fig. 8C)**. Metadata analysis did not identify any associated clinical or treatment-related factors (e.g., age, neoadjuvant therapy, hormonal treatment) that could account for the angiogenic EC enrichment. While angiogenic and tip-like endothelial phenotypes have been reported within tumor microenvironments across various cancers, to our knowledge, this study provides the first evidence of a systemic paraneoplastic effect in breast cancer, influencing endothelium in a normal tissue.

These results reveal a previously unrecognized systemic paraneoplastic influence of breast tumors on normal contralateral breast endothelial cells, with both contralateral and cancerous breast tissues showing markedly elevated angiogenic EC proportions compared to healthy controls. Together, these findings suggest that angiogenic ECs enrichment may serve as a biomarker of tumor-driven vascular remodeling and warrant further investigation into its potential diagnostic and therapeutic implications.

## Discussion

In this study, we characterized the heterogeneity and traced the origin of 597 CECs from 287 individuals using single-cell RNA sequencing of peripheral blood CD34⁺ cells. The scale of our cohort, together with comprehensive human endothelial cell atlases generated from the Tabula Sapiens dataset and 17 additional scRNA-seq datasets encompassing 212,144 endothelial cells from 23 tissues, enabled us, for the first time, to demonstrate the rarity of CECs and to apply a quantitative framework tracing their diverse tissue and vascular origins.

We constructed a healthy human endothelial reference manifold, forming the foundation for a quantitative scoring system based on validated literature-derived EC markers (Fig. 1). This framework enabled the distinction of major endothelial subtypes, including lymphatic, arterial, venous, capillary, and angiogenic, and uncovers intermediate transcriptional types. Projecting tissue profiles onto the manifold revealed organ-specific signatures and enrichments of disease-relevant genes. Among these, we identified TNNT3 distinct expression in cardiac capillary endothelial cells, previously reported to be expressed only in extra-cardiac smooth and skeletal muscle, and validated its expression by immunohistochemistry.^70^ This framework can be further applied to other datasets, providing robust, standardized comparison of endothelial types, states, and tissue origins across datasets and atlases. Beyond classification, it permits systematic analysis and cross-comparison of previously uncharacterized and disease-associated endothelial cells, as demonstrated by the precise mapping of CEC diversity. Together, these capabilities establish a platform for resolving endothelial programs and tissue-specific alterations in health and disease.

Our findings show that CEC transcriptional profiles span arterial, venous, capillary, and intermediate subtypes, while tissue-level profiling detected CECs with signatures from heart, liver, brain, and skeletal muscle (Fig. 2). This indicates multiple blood vascular populations and tissue origins, challenging earlier reports that a major subset of these cells represents a distinct progenitor population derived from the bone marrow. ^71,72^ In contrast to prior studies, our analysis establishes a rigorous, objective framework for quantifying CECs, providing the most accurate estimates of their absolute proportions to date. This approach reveals that CECs are far rarer than some previously proposed, resolving the long-standing debate over their existence and prevalence. ^43,73^ Their scarcity also clarifies that while CECs hold promise as disease biomarkers, their scarcity presents a major obstacle to their reliable use in studying disease mechanisms, given current technological resolution.

Cultured CECs exhibit marked phenotypic plasticity, undergoing dedifferentiation toward a progenitor-like state *in-vitro*, yet retaining the capacity to re-differentiate into mature endothelial phenotypes in response to external signaling cues (Fig. 3 and 4). Over three and six weeks of culture, CECs showed reduced mature vascular-type scores compared to *in-vivo* ECs, accompanied by progressive loss of *CD34* and reciprocal gain of *PROCR* over time, consistent with dedifferentiation toward a non-mature progenitor-like endothelial state. Within this overall trend, we observed subpopulations resembling *in-vivo* counterparts, including high-shear pressure–like ECs, low-shear pressure–like ECs, and angiogenic EC subsets. Altering external cues via Notch signaling inhibition or activation restored *CD34* expression and reduced *PROCR*, reversing the dedifferentiation seen in culture. Notch activation further generated cells with high angiogenic and capillary scores, comparable to mature counterparts in the reference Tabula Sapiens EC atlas. Together, these results demonstrate the capacity of cultured CECs to re-differentiate and establish an *in vitro* angiogenic model using cultured CECs.

To illustrate the utility of our reference manifold and quantitative angiogenic profile in detecting disease-associated endothelial programs, we analyzed a breast cancer EC scRNA-seq dataset, identifying a high angiogenic EC fraction, consistent with a tip-like EC phenotype reported in cancer (Fig. 5). ^63,68^ Unexpectedly, analysis of a normal breast tissue dataset also revealed an elevated angiogenic EC proportion in the contralateral “normal” breast tissue from breast cancer patients compared to healthy donor breast tissue (Fig. 5). This enrichment is independent of clinical or treatment variables and represents, to our knowledge, the first evidence of a systemic paraneoplastic effect on ECs in distant normal tissue. These findings warrant validation in additional datasets and call for research into its potential for early detection and therapeutic intervention.

To conclude, our comprehensive single-cell human endothelial atlases trace the diverse tissue and vascular origins of CECs and provide a quantitative framework for defining endothelial types and states across datasets. Applying this framework to breast cancer revealed angiogenic programs in tumors and a previously unrecognized paraneoplastic angiogenic program in contralateral normal breast tissue, demonstrating its value for detecting disease-associated endothelial changes. Finally, the bidirectional differentiation plasticity of CECs, dedifferentiating in culture yet re-differentiating into mature states with targeted signaling, positions them as a tractable model for dissecting tumor-driven angiogenic programs. This intrinsic plasticity also suggests *in vivo* potential that warrants exploration in future studies.

## Methods

### Tabula Sapiens EC model production

#### Data source and preprocessing

We analyzed publicly available scRNA-seq data from the Tabula Sapiens consortium, a comprehensive human atlas generated across multiple tissues and organs. ^27^ We included only cells produced using the 10x Genomics Chromium Single Cell 3’ Reagent Kits V3 or V3.1 to prevent technological biases between samples. Unique molecular identifier (UMI) counts and gene expression matrices were downloaded directly from the consortium’s repository.

#### Endothelial cell filtering and metacell model construction

Cells with UMI counts below 1,000 or above 12,000 and those with >0.2 mitochondrial UMI fraction were removed. A metacell model was built from the remaining cells. Endothelial metacells were defined by *CDH5* UMI fraction > 10^-4 or 10^-4.5 in some datasets to ensure representation and *PECAM1* UMI fractions >10^-4. Non-endothelial metacells were excluded if they expressed known pericyte (*PDGFRB*), smooth muscle (*MYH11*), epithelial (*S100A7*), lymphocyte (*LST1*), megakaryocyte (*PF4, PPBP*), club cell (*CYP2B7P*), basal cell (*S100A2*, *LAMB3*), mesenchymal (*COL6A3*), fibroblast (*THY1*), or hematopoietic-related (*CD14, HBB*) markers. Metacells with UMI fractions >10^-4 for any marker or >10^-3 for HBB were removed. The final endothelial cell set comprised cells from metacells passing these filters and was used for downstream analysis.

#### Data aggregation and metacell construction

A metacell model was generated from Tabula Sapiens-derived endothelial cells using the Metacell2 framework, with a target of 320K UMIs per metacell. Lateral genes, including cell cycle, heat shock, and stress markers such as *FOS* and *JUN*, were defined, and noisy genes were identified as bursty subsets of the lateral gene set. The model aggregated 20,891 cells into 582 metacells, with 78 outliers.

#### Vascular type annotation and marker curation

##### Literature-based scoring of vascular type endothelial markers

Established endothelial cell markers distinguishing vascular types were curated from peer-reviewed literature and categorized into blood vascular (arterial, venous, capillary) and lymphatic vessel endothelial cells (LVEC), as detailed in **Supplementary Table 1**. Each vascular type category was assessed based on how well its markers distinguished the corresponding metacells: LVEC, venous, arterial, and capillary ECs. This assessment involved validating markers identified from published literature by cross-referencing them with the original vascular type tissue annotations provided in the dataset, where available. Markers with strong discriminatory power were included in each vascular type score to ensure the correct classification of endothelial cell subsets across vascular types.

##### Calculating and validating vascular type scores

Vascular type scores were defined as the logarithm of the total expression (sum across markers) of the corresponding vascular genes, computed per metacell and individual cell. To validate the scores, the Kruskal–Wallis H-test and the Mann–Whitney U test were performed on the Tabula Sapiens ECs with vascular type annotation. The lymphatic score effectively distinguished lymphatic cells from non-lymphatic cells (p < 0.05), while the arterial, venous, and capillary EC scores were validated by comparing the respective cell types to others (p < 0.05).

Additionally, we calculated the following scores: an angiogenic score, which includes the genes ESM1, PXDN, PGF, ANGPTL2, and CXCR4; an HSP score, which includes the genes PROCR, SULF1, ELN, and BGN; and a BVEC score, which comprises genes that are anti-correlated with the LVEC marker genes. These latter scores were not validated due to the absence of a corresponding annotation in the Tabula Sapiens dataset and the fact that BVEC cells were identified by other scores.

##### Metacell annotation using validated vascular type scores

After constructing these literature-based scores, each metacell was assigned a score for each vascular type using the validated scoring system. The metacell was then annotated to the vascular type, corresponding to the highest score it received. For cases where the highest score was intermediate between arterial ECs and capillary ECs, the annotation was designated as "arterial-capillary," and when it was between capillary and venous, the annotation was designated as "capillary-venous." Angiogenic and HSP metacells were identified by their high angiogenic or HSP EC scores, respectively. When these scores overlapped with a high vascular type score, the annotation reflected this combination.

##### Score-based extrapolation of novel markers

To identify novel markers for each vascular type and angiogenic ECs, we analyzed genes with correlated expression (Pearson correlation coefficient >0.6) and differentially expressed genes in specific vascular type metacells. Iterative refinement excluded genes with broad expression across endothelial populations.

#### Cell cycle analysis

To capture the cell cycle phase of different metacells, we aggregated the expression of genes that are expressed in M-phase to one gene module, and the genes that are expressed in S-phase to another gene module. The score for each cell cycle phase was calculated as the logarithm of the mean expression of its corresponding gene module.

### Tissue endothelial cell models production

#### Data source and preprocessing

We utilized publicly available scRNA-seq datasets from diverse tissue types sampled from healthy cohorts, as detailed in **Supplementary Table 2**. To minimize potential batch effects, we included only studies conducted with 10x Genomics 3’ v3 or v3.1 sequencing technology. The raw sequencing data, including UMI counts and gene expression matrices, were directly downloaded from their respective repositories. Each dataset was processed individually to construct metacell models. Quality control steps and endothelial cells were filtered using the same criteria as in the Tabula Sapiens-based model to ensure consistency. A subset of datasets was re-processed, including a re-run of Cell Ranger (version 6.1.2) (**Supplementary Table 2**).

#### Vascular type classification and annotation of individual tissue models

We utilized the projection pipeline with default parameters to project the Tabula model onto each tissue model. ^74^ This approach provided basic annotations for each tissue metacell and facilitated the comparison of gene expression between each model and the Tabula model. Each metacell’s predicted vascular type classification was further manually reviewed and adjusted to ensure alignment with the vascular type scoring system. This approach ensured that each endothelial metacell was assigned to the vascular type most consistent with its transcriptional profile and tissue of origin.

#### Identifying tissue-specific markers

To identify genes specifically enriched in a specific tissue, we performed differential expression analysis between metacells from that tissue and all other relevant metacells. Because gene representation was not consistent across models, for tissues derived from the Tabula Sapiens EC model, comparisons were restricted to other metacells within the Tabula Sapiens EC model to ensure consistency in representation. For tissues derived from other datasets, comparisons were made against metacells from other tissue-specific models. A gene was considered a candidate to be significantly enriched if it showed (1) a relative log expression ratio ≥ 1, and (2) a statistically significant difference in expression distribution between the groups (χ² test, *p* < 0.05). For analyses stratified by vascular subtype, comparisons were performed within metacells of the same vascular identity.

Markers for specific tissues were selected following the exclusion of lateral and noisy genes (see ‘Data Aggregation and Metacell Construction’ above). The top 20 expressed genes for each vascular type within each tissue, as identified by the differential expression analysis, were examined. Marker gene selection was based on two main criteria: (1) widespread expression within the specific tissue (>20% of the tissue metacells showed high expression with a UMIs fraction of at least 4e-5), and (2) tissue specificity (a high-expression ratio >2.5 between the target tissue and all others). A metacell model was constructed using the full Tabula Sapiens dataset to screen for potential ambient noise. For each gene, its expression in ECs was compared to that of non-ECs within the same tissue. Genes with expression levels in non-ECs equivalent to or higher than those in ECs were designated as possible ambient noise and subsequently disregarded as tissue-specific EC markers.

#### Immunohistochemistry staining for Tropoin T3 expression

##### Tissue collection

Cardiac biopsy specimens were obtained from a 15-year-old female patient diagnosed with tetralogy of Fallot who was undergoing open-heart surgery repair. Immediately after excision, the tissue was kept on ice and then transferred to 4 mL of 4% paraformaldehyde (PFA) for fixation. Samples were then incubated for 24 hours under constant agitation in a cold, dark room.

##### Immunofluorescence staining

Paraffin-embedded heart tissues were sectioned and deparaffinized. Antigen retrieval was performed using 10 mM citric acid (pH 6.0). Sections were then permeabilized with 0.2% Triton X-100 in PBS for 5 min, followed by blocking with PBS containing 5% horse serum and 3% bovine serum albumin for 1 hour at room temperature. After blocking, sections were incubated overnight at 4°C with the following primary antibodies: anti-human cardiac Troponin T2 (CT3 hybridoma product, DSHB by Lin, J.J.-C., 1:10), anti-human CD31 (Abcam, ab28364), and anti-human Troponin T3 (Abcam, ab175058). On the following day, sections were washed with PBS and subsequently incubated with the appropriate secondary antibody (1:200, Abcam) for 30 minutes at room temperature. Nuclei were observed with DAPI (4,6-diamidino-2-phenylindole dihydrochloride) (SIGMA D9542, 5ug/ml).

Images were obtained using a Nikon Eclipse Ti2 fluorescent microscope equipped with Nikon’s NIS-Elements imaging software.

### CEC model productions

#### Sample Collection and Preparation

Peripheral blood samples were collected from 121 healthy volunteers 58 males and 63 females) aged 26 to 95 and 166 patients diagnosed with hematological malignancies, including MDS and MPN 99 males and 67 females) aged 22 to 96. Written informed consent allowing sequencing data generation (including genotyping panels) was obtained from all participants. All relevant ethical regulations were followed per the Declaration of Helsinki, under an ethically approved protocol by the Weizmann Institute of Science ethics committee (IRB protocol 283-1).

Patients’ basic demographic and clinical data are summarized in **Supplementary Table 3**. A 50 mL blood sample was drawn from each volunteer into lithium-heparin tubes and processed within 4 hours post-collection to preserve RNA integrity and the viability of cCD34+ cells. A small quantity of blood (1ml) was used for DNA sequencing, while the remaining blood was used for Peripheral blood mononuclear cells (PBMCs). PBMCs were isolated using Ficoll density gradient centrifugation in Lymphoprep-filled SepMate™ tubes (StemCell Technologies). CD34+ cell enrichment was then performed using the EasySep™ Human CD34 Positive Selection Kit II (STEMCELL Technologies), according to the manufacturer’s instructions. This method consistently achieved an enrichment purity of 50% to 95%. Flow cytometry verification was conducted using antibodies specific to CD34 (APC-Cy7-conjugated, BioLegend) and CD45 (Brilliant Violet 510-conjugated, BioLegend), confirming the characteristic of a live CD34+CD45int profile of CD34+ cells.

#### ScRNA-seq library preparation and processing of circulating CD34⁺ cells

The complete methodology for single-cell RNA sequencing (scRNA-seq) library preparation, sequencing, and computational analysis was performed as previously detailed in *Furer, Rappoport, Milman,* et al. (2025). ^42^ In summary, libraries were generated from circulating CD34+ cells using the 10x Genomics Chromium 3’ v3.1 kit within five hours of blood collection. Libraries were sequenced on both Illumina and Ultima Genomics platforms, with a comparative analysis confirming high similarity between the platforms. Following sequencing, we implemented the data processing and demultiplexing pipeline, as described in *Furer, Rappoport, Milman*, et al. (2025). Raw data were processed with Cell Ranger, and cells were filtered based on quality control metrics. To assign each cell to its donor of origin from the pooled samples, we used our established genotype-based strategy that integrates Vireo, Souporcell, and a metacell-based model for robust doublet identification and removal. ^75,76^ For a comprehensive description of the entire protocol, including specific reagents, sequencing parameters, and the full computational workflow, please refer to the original study. ^42^

#### Filtering for endothelial cells and final metacell model generation

Endothelial cells from CD34+ circulating scRNA-seq were identified by constructing a metacell model post-doublet filtering, using lateral and noise gene sets from the healthy endothelial cell models. Potential endothelial metacells were initially defined as those with a CDH5 UMI fraction >10^-4.5. To eliminate non-endothelial cell contaminants, a second metacell model was constructed using only the cells from this potential set. Metacells were classified as endothelial if both CDH5 and PECAM1 UMI fractions exceeded 10^-4. The final metacell model included cells passing both filters, with a target of 30 cells and 100K UMIs per metacell.

#### Comparing tissue markers in circulating endothelial cells to reference endothelial cells

For metacells comparison, tissue-specific marker genes were chosen from the tissue gene modules specified before. CEC metacells were compared only to the Tabula Sapiens EC metacells model, taking only metacells matching the tissue of comparison (as described above).

#### Matching tissue of origin to circulating endothelial cells

Circulating endothelial cells were downsampled to 1024 UMIs, removing 181 cells (30.32%), to minimize variability due to sampling. For each cell, we computed the total UMI count for each tissue-specific gene set and calculated its log-transformed fraction, using an epsilon of 1/1024. Cells were considered a potential match if their tissue module expression exceeded −7, indicating at least 8 UMIs per cell for the relevant tissue module, and are shown in the main figures. Cells processed without the downscaled process can be found in **Extended Data Fig. 5F**.

### Culturing of circulating endothelial cells and model production

#### Culture of circulating endothelial cells

45ml of peripheral blood obtained from 13 healthy adult donors under institutional ethical guidelines approved by the Weizmann Institute of Science ethics committee (IRB protocol 2531-1) and following informed consent procedures. PBMCs were isolated, with a small quantity of blood (1ml) used for DNA sequencing, as described previously.

Cultured circulating endothelial cell (CultECs) growth medium was prepared by supplementing Endothelial Basal Medium-2 (EBM-2; Cat. no. CC-3156, Lonza) with the EGM™-2 BulletKit™ SingleQuots™ (Cat. no. CC-4176, Lonza) according to the manufacturer’s protocol. The medium was further supplemented with fetal bovine serum (FBS, Cat. no. 26140079, Gibco) to reach 18% (v/v). The complete medium was sterile filtered using a 0.2 µm pore filter and stored at 4 °C for up to two weeks.

A Type I collagen solution was prepared for flask coating. Lyophilized CellAdhere™ Type I Collagen (Cat. no. 07005, STEMCELL Technologies) was reconstituted by adding 10 mL of 0.01 N HCl to 15 mg of collagen to produce a 3 mg/mL stock solution in a cell culture hood. The stock was further diluted by adding 500 µL of the 3 mg/mL collagen solution to 29.5 mL of 0.01 N HCl, yielding a final concentration of 50 µg/mL. Larger volumes were prepared by scaling appropriately (e.g., 750 µL of stock in 44.25 mL of 0.01 N HCl). The collagen solution was then applied to the cultureware surface at a final concentration of 5 µg/cm². For example, 2.5 mL of the 50 µg/mL collagen solution was applied to a T-25 flask (providing 125 µg of collagen per flask). For multi-well plates, the volume was scaled appropriately to maintain the same surface coverage. Following collagen application, the coated cultureware was incubated at RT for 1 h. Following incubation, the excess collagen solution was aspirated, and the flasks were washed twice with PBS to remove residual acid (e.g., 5 ml of PBS in a T-25 flask). If PBMCs were not immediately ready for seeding, the CultECs growth medium was added to each flask to prevent the collagen coating from drying until cell plating (e.g., 2 ml in a T-25 flask).

PBMCs were plated immediately after isolation on the collagen-coated cultureware. Following centrifugation and washing, the PBMC pellet was resuspended in 1 mL of CultECs growth medium and pipetted thoroughly to achieve a single-cell suspension. Each donor’s PBMCs were cultured and eventually plated individually. The total volume was adjusted with the CultECs growth medium to achieve the desired plating density based on the cultureware used. For example, for a T-75 flask, the entire cell suspension, containing approximately 40 million PBMCs from 40 mL of blood, was seeded with a total volume of 15 mL completed with the CultEC medium. Cultures were incubated at 37 °C in a humidified atmosphere with 5% CO₂. The initial culture was designated as passage 0.

The culture medium was replaced every two days by aspirating the old medium and adding fresh, pre-warmed (37 °C) CultECs growth medium. Once a week, medium changes were extended to three days if necessary.

Cultures were observed daily under a phase-contrast microscope for cell attachment and proliferation. Endothelial cell colonies, characterized by cobblestone morphology, typically appeared between 14 and 21 days.

#### Passage of cultured circulating endothelial cells

For T-25 flasks, cells were rinsed twice with 3 mL of DPBS. Subsequently, 2 mL of pre-warmed (37 °C) 1X trypsin-EDTA (Cat. no. 25200056, Gibco) was added, and the flasks were incubated at 37 °C for 5 min. After incubation, trypsin was neutralized with 5 mL of CultEC growth medium containing 18% FBS, and the cells were brought into suspension with repeated pipetting. The cell suspension was centrifuged at 300 × g for 5 min, the supernatant was discarded, and the pellet was resuspended in 5 mL of fresh CultECs growth medium. The entire cell suspension was plated into a new T-25 flask, representing passage 1 (P1). For other surface-area flasks or plates, volumes were scaled accordingly. No collagen coating was required for subsequent passages. Medium changes were continued as described.

#### Matrigel assay of cultured circulating endothelial cells

CultECs were plated on growth factor–reduced Matrigel (Corning, 354230) following the manufacturer’s protocol. For each well of a 24-well plate, 289 µL of Matrigel was dispensed, and then incubated at 37°C for 30–60 minutes to allow gel polymerization. Following Matrigel solidification, 300 µL of cell suspension containing 1.2 × 10⁵ CECs was added to each well. The plates were then incubated at 37°C in a humidified atmosphere with 5% CO₂ for 16–18 hours. Morphological assessments and network formation were evaluated every 6 hours.

#### Differentiation assays on cultured circulating endothelial cells

To assess cell plasticity and differentiation potential, CultECs were cultured for five days under specific conditions by adjusting a differentiation protocol by Ang et al. 2022. ^60^ Two differential conditions were applied: (1) a non-direct Notch activation, using CultEC growth medium excluding the kit-supplied VEGF, supplemented with only 2% FBS, vascular endothelial growth factor (VEGF; 100 ng/mL, Cat. no. 130-109-383, Miltenyi Biotec), and Activin A (25 ng/mL, Cat. no. 130-115-008, Miltenyi Biotec). ^77,78^ (2) A direct Notch inhibition, using CultEC growth medium supplemented with 10% FBS and the γ-secretase inhibitor RO4929097 (1 µM, Cat. no. sc-364602, Santa Cruz Biotechnology). ^79^ The media were replaced daily during the five-day duration of the differentiation experiment. Cells were monitored daily under a phase-contrast microscope.

#### Phase contrast image capture of cultured endothelial cells

Phase-contrast images were acquired using an Olympus microscope equipped with an XM10 monochrome CCD camera. Image capture and initial processing were performed using Olympus Soft Imaging Solutions software.

#### Collection and scRNA-seq of cultured endothelial cells

Cells were enzymatically dissociated into a single-cell suspension, as described above under "Passage of CultECs". Following cell suspension, cells were washed twice by centrifuging at 300 × g for 5 min and resuspended in 1 mL of CultECs growth medium. Libraries for cultured endothelial cells were generated using the 10x Genomics Chromium Next Gem Single Cell 3’ Reagent Kits (v3.1 and v4) according to the manufacturer’s instructions. Approximately 30,000 cells were loaded per Chromium Chip. Libraries were sequenced on Illumina NovaSeq 6000 or NovaSeq X platforms. Demultiplexing, data processing, and metacell model generation were performed as previously described. For the v4 libraries, a newer version of cell-ranger (8.0.1) was used for processing the data.

#### Annotation of cultured circulating endothelial cells

CultECs were annotated based on gene expression profiles. Five primary subpopulations were identified, along with two additional subsets exhibiting an angiogenic-EC state signature, described previously.

Two main groups were annotated based on their similarity to ECs from the reference Tabula Sapiens EC model: (1) High-shear pressure–like endothelial cells (HSP-like ECs). This group was defined by a high expression of genes associated with high-shear pressure, including *PROCR, SULF1, ELN*, and *BGN*. ^9,33^ Additional markers, such as extracellular matrix components (*LTBP2, MMP2), NRG1, ART4*, and arterial markers (*GJA4, LTBP4*), observed in this population, also supported this classification. A subgroup within this population, which also showed elevated angiogenic scores, was designated HSP-like angiogenic-ECs. (2) Low-shear pressure–like endothelial cells (LSP-like ECs). This group was annotated based on its high expression of genes enriched in venous endothelial cells from the Tabula Sapiens atlas. This population was distinct from the HSP-like ECs, lacking high expression of their characteristic genes. Key markers for this group included *AQP1, C7, TFPI*, and *CCL14*. Finally, three additional subpopulations were characterized by the expression of stress-associated genes, such as *NQO1* and *HMGA1*. These groups were named according to their varying levels of HSP-like and LSP-like markers, indicating a spectrum of stress responses.

Cultured CECs undergoing differentiation assays were annotated based on their condition, using a majority vote of the cells in each metacell (i.e., >75%).

#### Differential expression analysis comparing cultured circulating endothelial cells undergoing differentiation assays

To identify genes enriched in a specific condition, we performed differential expression analysis between metacells from different conditions using the MCView Shiny tool (https://github.com/tanaylab/MCView). ^80^

### Breast tissue EC analysis

A metacell model of ECs from both healthy and cancerous breast tissue was built using publicly available data following the same methodology used for the Tabula Sapiens and other publicly available tissue datasets. We annotated the models by assigning endothelial type and state scores to each metacell, as described earlier. To ensure a representative sample size, comparing the fraction of angiogenic ECs was done only in patients with more than 50 and 500 cells. The statistical significance of this comparison was calculated using___the Mann-Whitney U test. Similarly, the distribution of EC types and states across patients was restricted to those with at least 50 and 500 cells.

### UMAP calculation

Each metacell model was projected onto a 2D UMAP using either the native metacell UMAP calculation functions or directly using the Python UMAP library, with the various scores for each metacell as input. ^81^ These included arterial, capillary, venous, LVEC, BVEC, and angioeginc-scores.

### UMAP and marker heatmap for healthy endothelial cell states

The marker heatmap and UMAP for the healthy endothelial cells were calculated similarly to those of each dataset. A combined presentation of the marker heatmap was done by combining the models into one.

### Computational and statistical tools

All analyses were performed using Python (version 3.12.4). Key libraries included Metacell2 (version 0.9.5) for metacell modeling. ^28^ Data and code used in this study are available upon reasonable request. All significant calculations comparing scores were done by the Kruskal–Wallis H-test and the Mann–Whitney U test unless stated otherwise.

### Data and code availability

All data and code generated in this study will be provided prior to publication.

## Supplementary Information

**Supplementary Table 1** – **Literature-validated endothelial cell markers used in our scoring system.** This table lists endothelial cell markers curated from the literature and validated for their ability to discriminate between endothelial cell types (arterial, venous, capillary, and lymphatic endothelial cells).

**Supplementary Table 2** – Datasets included in the construction of individual tissue-specific endothelial cell atlases.

**Supplementary Table 3 - Demographic characteristics of circulating CD34⁺ sample donors.** This table summarizes the sex, age, and hematological disease background of all participants from whom circulating endothelial cells were sampled.

**Supplementary Table 4** – Differential gene expression in cultured circulating endothelial cells under different Notch pathway alteration conditions.

---

**Extended Data Fig. 1.**
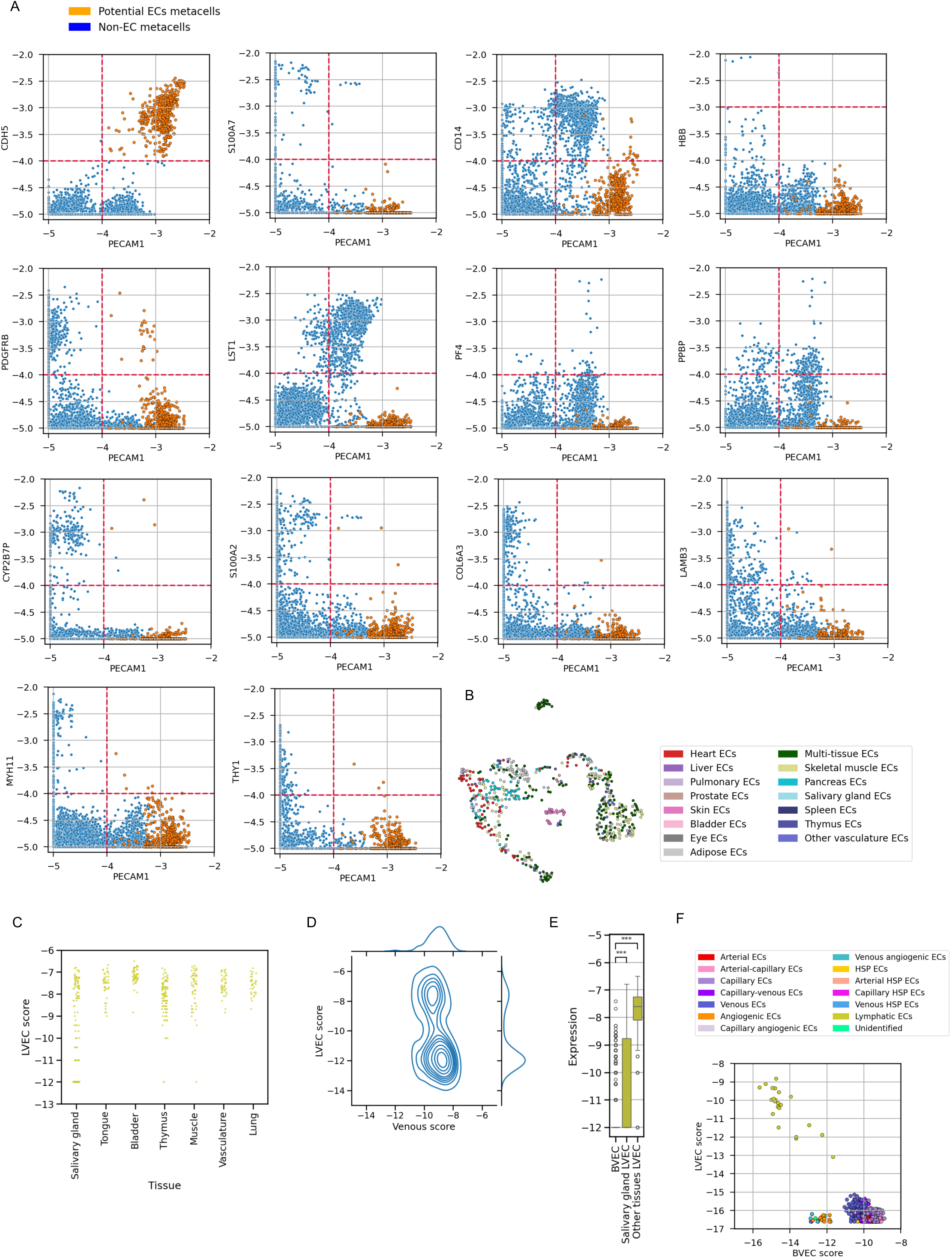
A. Scatter plots of gene expression applied to filter endothelial cells from the Tabula Sapiens dataset. Each dot represents a metacell. Orange dots represent metacells consistent with an endothelial cell identity that exhibit high expression of *CDH5* and *PECAM1*, and were filtered to exclude non-endothelial cells. B. Two-dimensional UMAP projection of the transcriptional manifold of Tabula Sapiens endothelial cells. Metacells containing at least 75% cells from the same tissue were annotated by the tissue, while a mixture of tissues was annotated as multi-tissue ECs. C. Lymphatic score per cell across tissues in the endothelial Tabula Sapiens atlas, based on original single-cell annotations. Each dot represents a single cell. D. Kernel density estimation (KDE) plot of lymphatic versus venous scores in salivary gland endothelial cells. The distribution reveals two main populations: one expressing both lymphatic and venous markers, and another enriched for venous signature only, suggesting misannotation of the latter as lymphatic. E. Lymphatic vessel endothelial cells (LVEC) and blood vessel endothelial cells (BVEC) score expression at the cell level for salivary gland lymphatic annotated cells. Each dot represents a cell. Despite potential misannotations, salivary gland LVECs show significantly higher LVEC scores compared to BVECs. F. Scatterplot showing BVEC versus LVEC score expression in the endothelial Tabula Sapiens model. Each dot represents a metacell.

**Extended Data Fig. 2.**
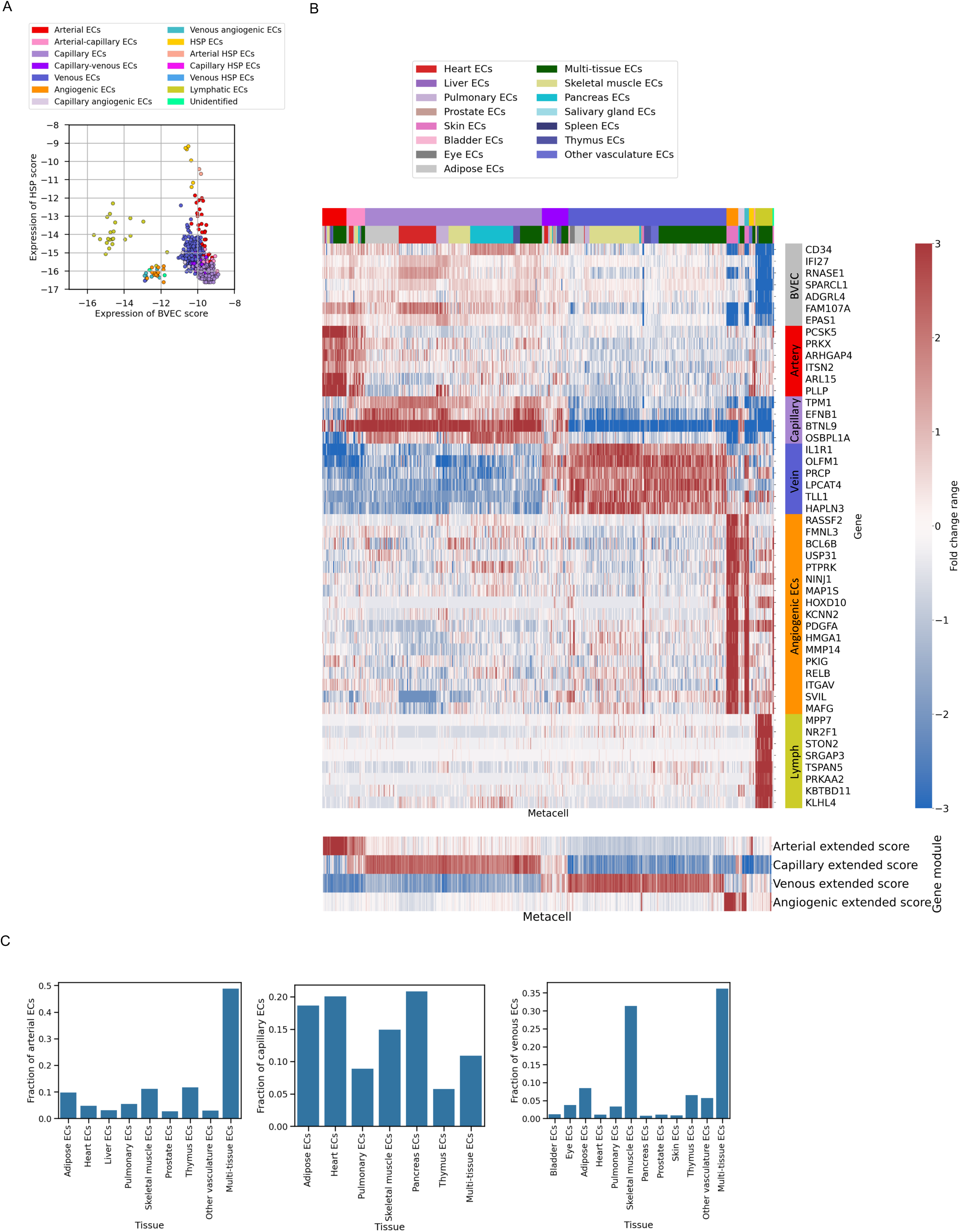
A. Scatter plot BVEC versus HSP score expression across metacells in the endothelial Tabula Sapiens model. Colors indicate endothelial type and state. Each dot represents a metacell. B. Heatmap of novel endothelial type markers across metacells in the Tabula Sapiens atlas. Genes were selected for high expression and correlation with endothelial type. The top bar indicates endothelial type (top row, color-coded as in Extended Data Fig. 2A) and tissue of origin (bottom row, color-code provided separately). The lower panel shows extended type scores, computing both literature-based and novel markers. C. Bar plot showing the fraction of arterial, capillary, and venous endothelial cells across tissue.

**Extended Data Fig. 3.**
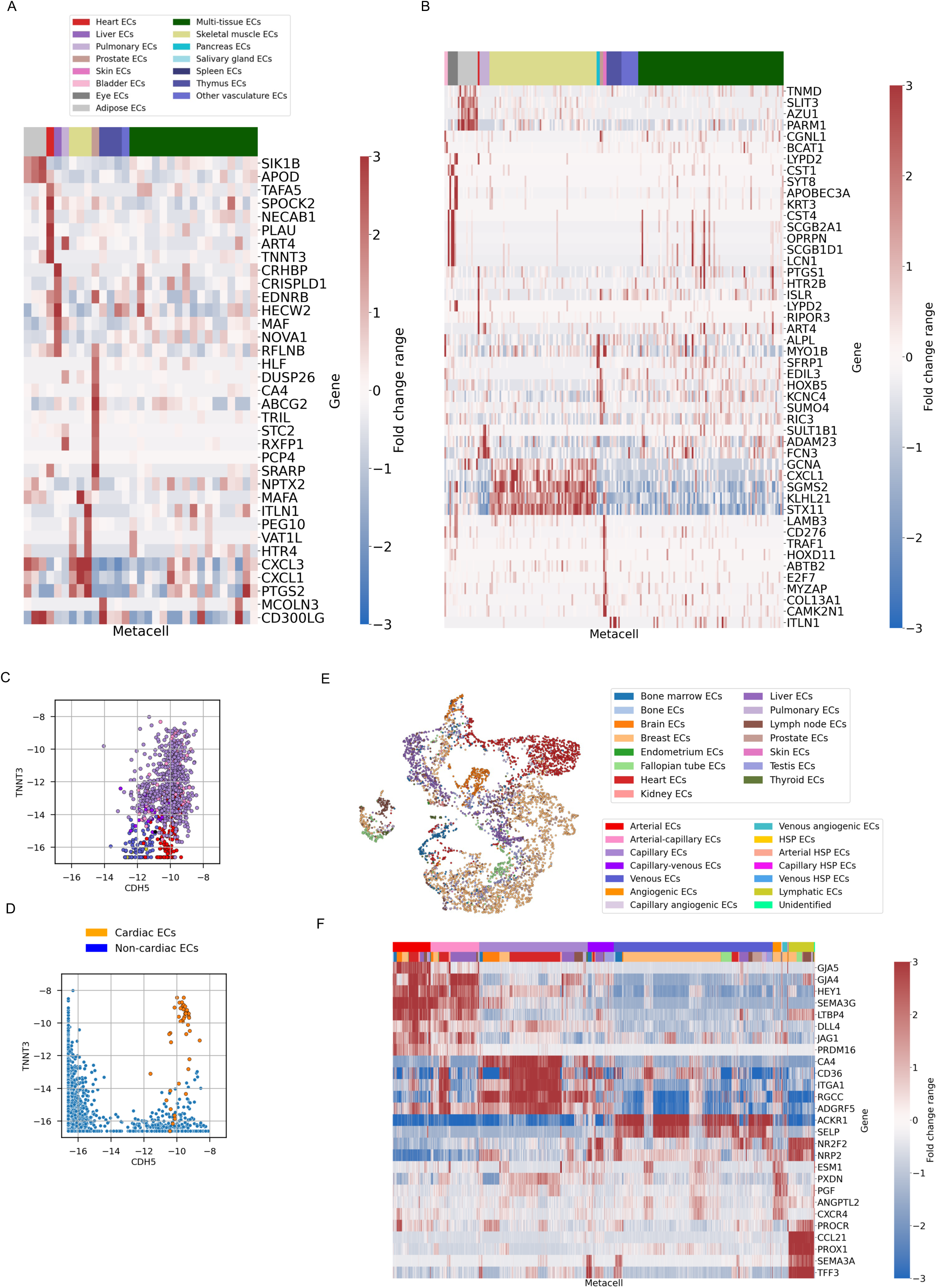
A. Heatmap of differentially expressed tissue-specific marker genes across arterial endothelial metacells from the endothelial Tabula Sapiens atlas. The top bar indicates the tissue of origin. B. Heatmap of differentially expressed tissue-specific marker genes across venous endothelial metacells from the endothelial Tabula Sapiens atlas. The top bar indicates the tissue of origin. C. Scatter plot of *CDH5* versus *TNNT3* expression in cardiac endothelial cells from Kanemaru, K., Cranley, J., Muraro, D. *et al*. High *TNNT3* expression is observed in the capillary cardiac endothelial cells. D. Scatter plot of *CDH5* versus *TNNT3* expression across all metacells in the full Tabula Sapiens atlas, including cardiomyocytes. Each dot represents a metacell. Orange dots represent cardiac endothelial metacells. E. Two-dimensional UMAP projection of the transcriptional manifold of endothelial cells derived from the tissue atlases. F. Heatmap of literature-based vascular-type endothelial markers across metacells in the tissue atlases. The top annotation indicates endothelial type (top bar) and tissue of origin (lower bar).

**Extended Data Fig. 4.**
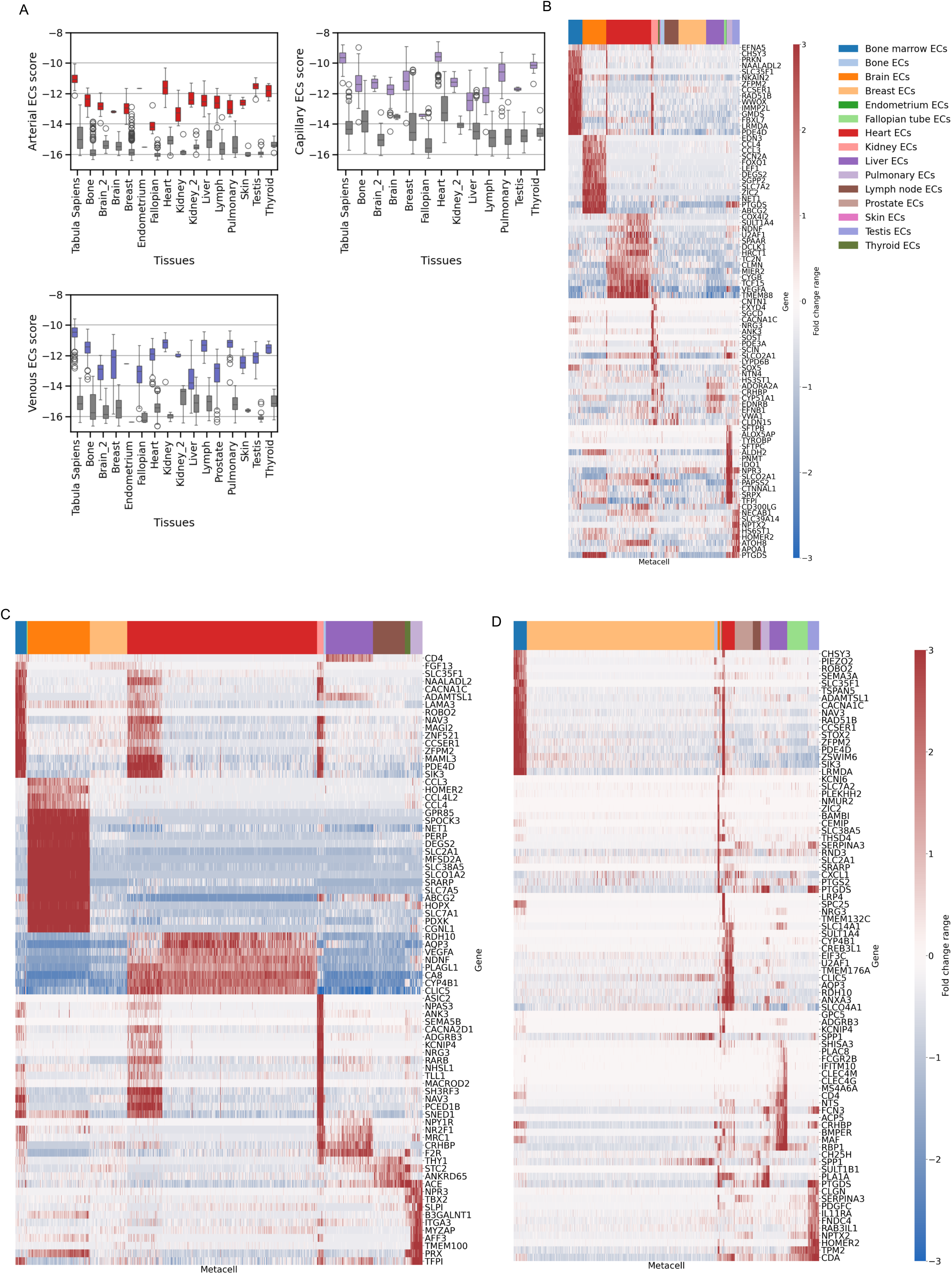
A. Expression of (clockwise) arterial, capillary, and venous endothelial type scores across tissues and models. Each dot represents a metacell. The boxplot color corresponds to the scored endothelial type. B. Heatmap of differentially expressed tissue-specific marker genes across arterial endothelial metacells from the tissue atlases. The top bar indicates the tissue of origin. C. Heatmap of differentially expressed tissue-specific marker genes across capillary endothelial metacells from the tissue atlases. The top bar indicates the tissue of origin. D. Heatmap of differentially expressed tissue-specific marker genes across venous endothelial metacells from the tissue atlases. The top bar indicates the tissue of origin.

**Extended Data Fig. 5.**
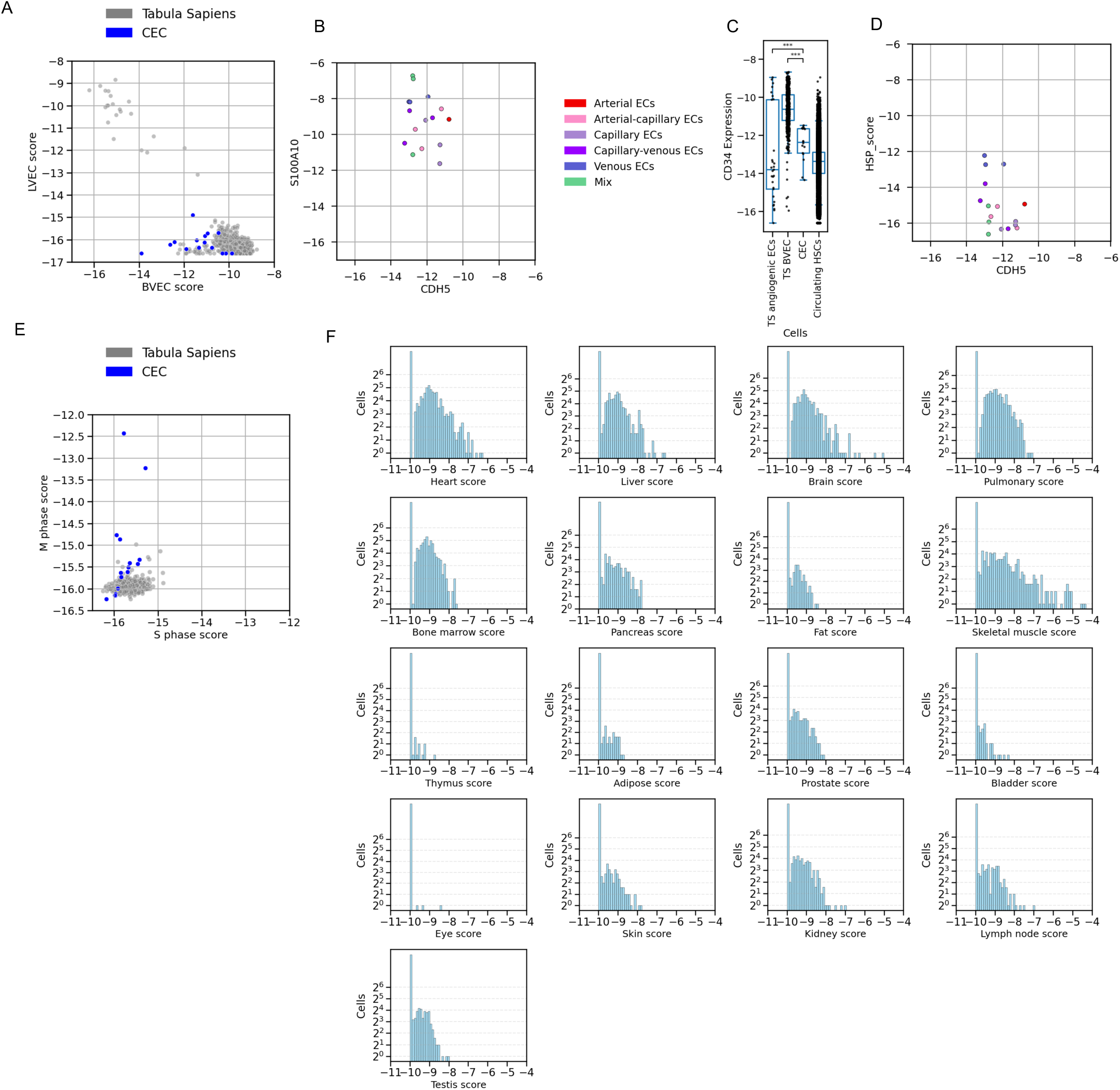
A. Scatter plot of blood vascular endothelial cells (BVEC) versus lymphatic vascular endothelial cells (LVEC) expression scores across metacells from the endothelial Tabula Sapiens model (gray dots) and from the circulating endothelial cell (CEC) model (blue dots), supporting a BVEC origin for CECs. B. Scatter plot of *CDH5* versus *S100A10* expression in CEC metacells. A subset of CECs co-express high levels of endothelial and stress-associated markers, suggesting stress-related activation. C. Boxplot of *CD34* expression across CECs, Tabula Sapiens (TS) angiogenic ECs, TS BVECs, and circulating CD34^+^ hematopoietic stem cells (HSCs). CD34⁺ peripheral blood sorting captures diverse endothelial populations, including angiogenic ECs. D. Scatter plot of *CDH5* versus heat shock protein (HSP) scores in circulating endothelial metacells. Low HSP expression indicates no distinct HSP-positive cell population. E. Scatter plot of cell cycle scores across CEC metacells, showing two metacells with high M-phase scores, indicative of active cycling. F. Histograms of tissue-specific score expression across circulating endothelial cells. Each histogram shows a different tissue with a subset of cells exhibiting elevated tissue-specific marker expression, supporting that tissue origin.

**Extended Data Fig. 6.**
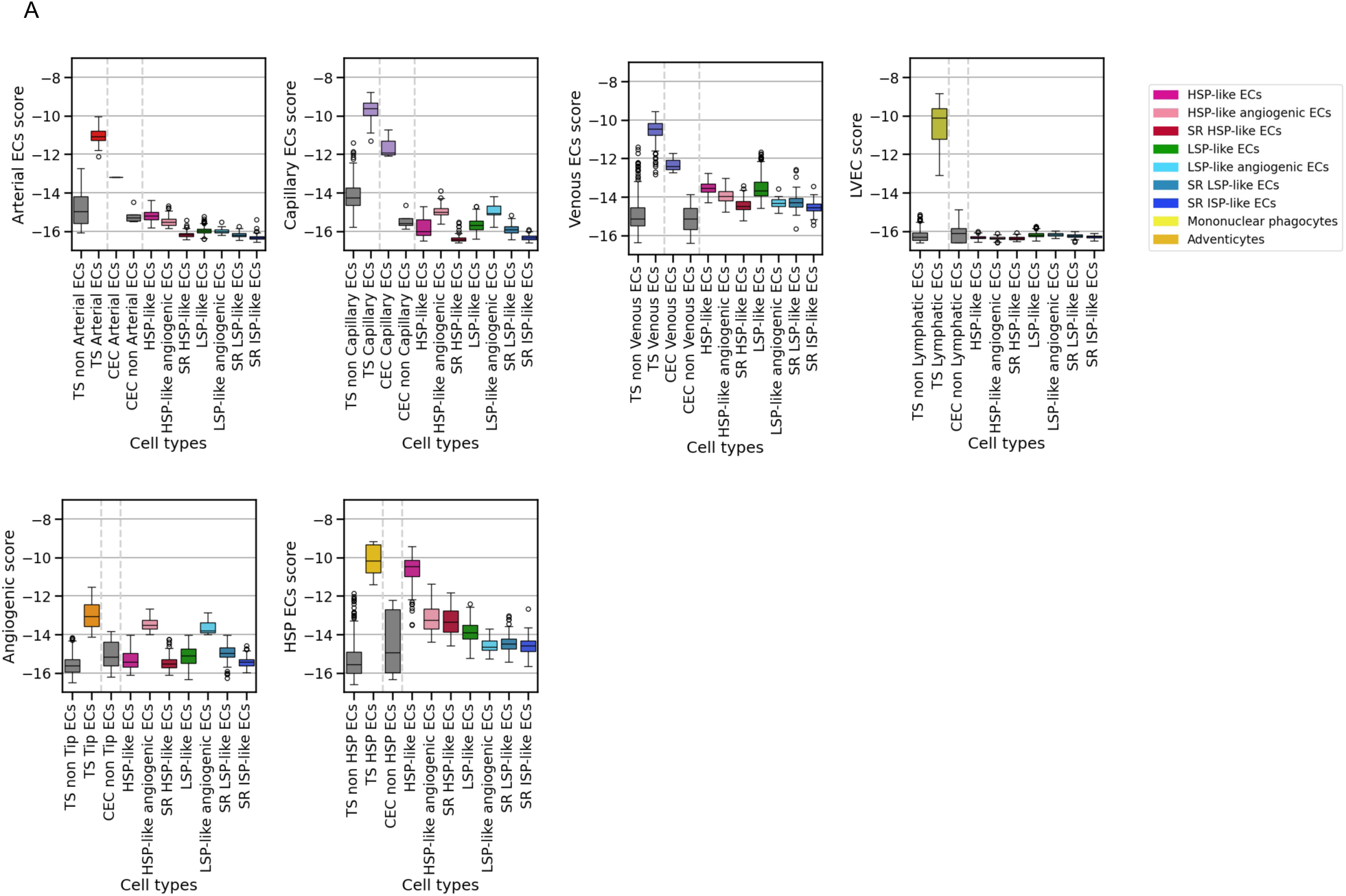
A. Boxplots of arterial, capillary, venous, lymphatic, and angiogenic endothelial scores across metacells from endothelial Tabula Sapiens (TS), circulating endothelial cells (CEC), and the different subgroups of cultured CEC, showing a progressive decrease of mature type endothelial cells from resident endothelial cells to CECs and through culture. Boxplot colors represent the corresponding endothelial type and state.

**Extended Data Fig. 7.**
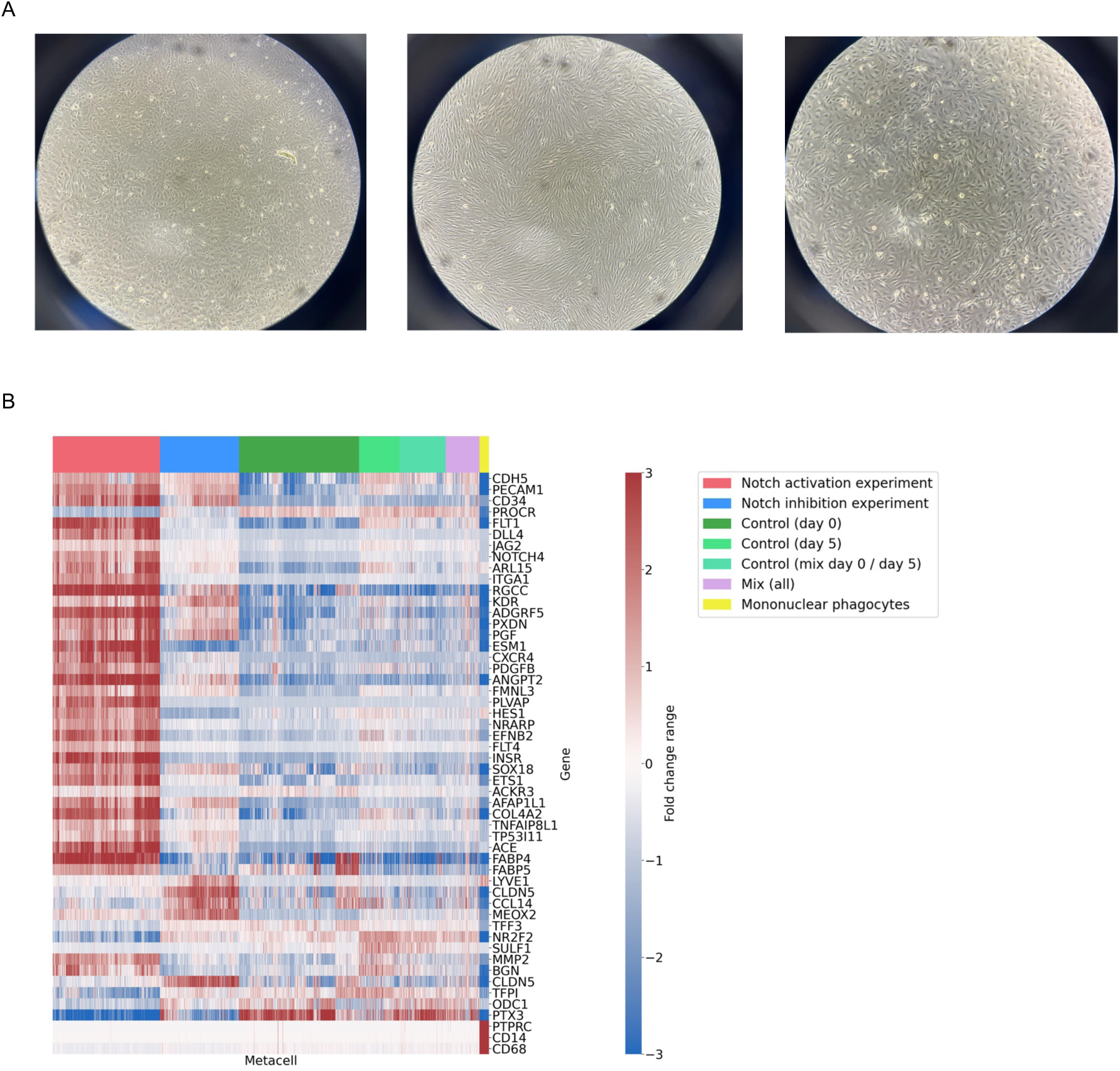
A. Phase-contrast microscopy images of cultured CECs under control conditions, Notch pathway activation, and Notch pathway inhibition (left to right). Morphological changes under Notch activation became evident after 12 hours. B. Heatmap of differentially expressed genes across cultured CEC metacells under Notch activation, Notch inhibition, control subpopulations (see Fig. 3), and the mononuclear phagocytic population. The top bar indicates the experimental condition; yellow represents mononuclear phagocytes.

**Extended Data Fig. 8.**
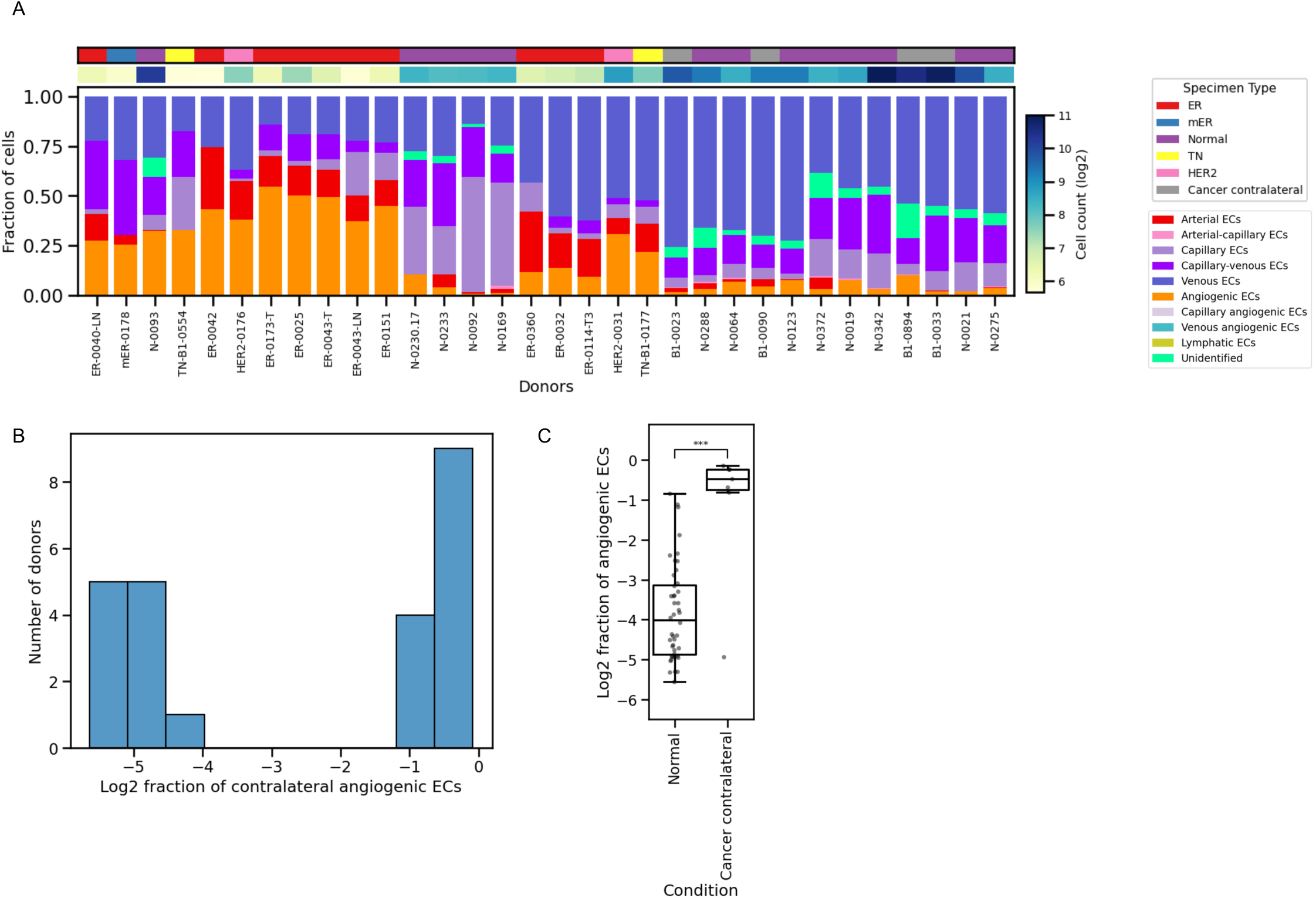
A. Composition of endothelial cell states at the single-patient level from Chen, Y., Pal, B., Lindeman, G.J. et al. Each column represents a patient, with colors indicating the relative proportion of annotated endothelial type and states. Only patients with at least 50 cells are shown. B. Histogram showing the distribution of contralateral breast tissue patient samples across angiogenic endothelial cell fractions. C. Boxplots of angiogenic cell fraction distributions across normal breast tissue of healthy individuals versus normal contralateral breast tissue of breast cancer patients (log scale). Patients with fewer than 500 cells were excluded.

